# Inhibition of sensory neuron driven acute, inflammatory, and neuropathic pain using a humanised chemogenetic system

**DOI:** 10.1101/2023.03.21.533690

**Authors:** Jimena Perez-Sanchez, Steven J. Middleton, Luke A. Pattison, Helen Hilton, Mosab Ali Awadelkareem, Sana R. Zuberi, Maria B. Renke, Huimin Hu, Xun Yang, Alex J. Clark, Ewan St, John Smith, David L. Bennett

**Author notes:** These authors contributed equally to this work (Jimena Perez-Sanchez, Steven J. Middleton).

## Abstract

Hyperexcitability in sensory neurons is known to underlie many of the maladaptive changes associated with persistent pain. Chemogenetics has shown promise as a means to suppress such excitability, yet chemogenetic approaches suitable for human applications are needed. PSAM^4^-GlyR is a modular system based on the human α7 nicotinic acetylcholine and glycine receptors, which responds to inert chemical ligands and the clinically-approved drug, varenicline. Here, we demonstrated the efficacy of this channel in silencing both mouse and human sensory neurons by the activation of large shunting conductances after agonist administration. Virally-mediated expression of PSAM^4^-GlyR in mouse sensory neurons produced behavioural hyposensitivity upon agonist administration, which was recovered upon agonist washout. Importantly, stable expression of the channel led to similar reversible behavioural effects even after 10 months of viral delivery. Mechanical and spontaneous pain readouts were also ameliorated by PSAM^4^-GlyR activation in acute and joint pain inflammation models. Furthermore, suppression of mechanical hypersensitivity generated by a spared nerve injury model of neuropathic pain was also observed upon activation of the channel. Effective silencing of behavioural hypersensitivity was reproduced in a human model of hyperexcitability and clinical pain: PSAM^4^-GlyR activation decreased the excitability of human induced pluripotent stem-cell-derived sensory neurons and spontaneous activity due to a gain of function Na_V_1.7 mutation causing inherited erythromelalgia. Our results demonstrate the contribution of sensory neuron hyperexcitability to neuropathic pain and the translational potential of an effective, stable and reversible human-based chemogenetic system for the treatment of pain.

## INTRODUCTION

Chronic pain remains a critical unmet clinical challenge. Approximately one in five adults suffers from chronic pain (*1*), which can arise as a consequence of tissue damage, disease or inflammation (*2, 3*). Current treatments have poor efficacy and tolerability (*4*); the devastating opioid crisis highlights the need for new, more effective, non-addictive treatments for this condition. However, the varied and complex pathophysiological mechanisms that underlie chronic pain have made this a particularly difficult task.

Increased activity in sensory neurons, which relay information from the periphery to the central nervous system (CNS), has been identified as a common driver for the induction and maintenance of inflammatory (*5*) and neuropathic pain following tissue injury (*6–8*). Sensory neurons rely on the generation of action potentials (APs) to encode such information. Increased AP spiking (hyperexcitability) is a hallmark of altered sensory neuron processing, observed as changes in firing pattern, bursting and the generation of spontaneous activity (*9–15*). Intrinsic excitability arises from the complex interaction of multiple ion channels (*16*), which means that countless molecular changes may be responsible for producing altered activity in different chronic pain conditions. Extrinsic factors such as immune mediators can also sensitise sensory neurons (*17*). So far, efforts to ascribe single molecular changes to altered activity have made headway to recognise potential therapeutic targets, but they have not necessarily been translated successfully into treatments (*4, 18, 19*). An alternative plan to guide future strategies is to surpass the molecular mechanisms that enhance sensory neuron excitability and directly suppress the activity in these neurons. Indeed, local anaesthetics that block the generation of APs in peripheral nerves transiently reduce both spontaneous and evoked pain in animal models of nerve injury, as well as in neuropathic and osteoarthritis pain patients (*8, 20–22*). Nevertheless, such non-specific approaches are not clinically viable, because systemic administration may give rise to motor, cardiac and CNS side-effects (*4*).

Chemogenetic approaches serve this purpose as they are able to modulate neuronal activity in defined neuronal populations, by expressing designer receptors that respond selectively to specific agonists which enables dose dependent control and reversibility (*23*). Ideally, these approaches should be compatible with human tissue to provide potential for gene therapy. Indeed, chemogenetic approaches have been used to suppress the activity of sensory neurons and pain signaling (*24–27*). However, several concerns have been raised regarding agonist viability of the most common chemogenetic tools, as well as their disruption of endogenous receptor function (*26, 28, 29*). Furthermore, chemogenetic strategies have never been used in human models of neuropathic pain as a means to suppress ectopic activity. The novel pharmacologically-selective actuator modules (PSAMs) are promising in this regard (*30, 31*). They are modular systems based on a modified human ɑ7 nicotinic acetylcholine (ACh) receptor ligand-binding domain, which does not bind to ACh, but ultrapotent pharmacologically-selective effector molecules (uPSEMs) and in the case of the most recent version PSAM^4^, the FDA-approved drug varenicline. This, in combination with the human glycine receptor pore domain, gives rise to PSAM^4^-GlyR, a chloride channel activated by selective agonists. This tool has been used to inhibit neuronal activity in the brain, although its inhibitory effect is highly dependent on the chloride gradient (*31, 32*). In this study, we used PSAM^4^-GlyR to silence the activity of mouse sensory neurons in acute, inflammatory and neuropathic pain conditions. Furthermore, to show its potential for human therapeutic application, we expressed PSAM^4^-GlyR in human sensory neurons, where we used it to silence neuronal activity, as well as ectopic activity in a human cellular model of neuropathic pain.

## RESULTS

### Generation of a new PSAM^4^-GlyR construct

We had to consider several factors when designing a PSAM^4^-GlyR construct for use in sensory neurons. We required a strong promotor, a red fluorescent tag, and a short, efficient linker. We modified the original construct containing PSAM^4^-GlyR (*31*) to account for these factors. We also ensured its size was reduced, suitable for packaging into adeno-associated virus (AAV) vectors (*33, 34*), in preparation for *in vivo* delivery to sensory neurons. After generating several candidates, we found the most promising construct to be pAAV-CAG-mCherry-tPT2A-PSAM4-GlyR (mCherry-T-PSAM^4^-GlyR) (see Methods). We assessed expression and function of our construct in HEK293t cells and used whole-cell patch clamp recordings to measure the induced membrane conductance following addition of varenicline, the clinically relevant agonist of PSAM^4^-GlyR (*31*). We confirmed that mCherry-T-PSAM^4^-GlyR was functional, and observed a large increase in membrane conductance following varenicline application, similar to the original unmodified construct (Fig. S1). We therefore selected mCherry-T-PSAM^4^-GlyR (and mCherry-alone to use as control) for all ensuing experiments (Fig. 1A).

**Fig. 1.**
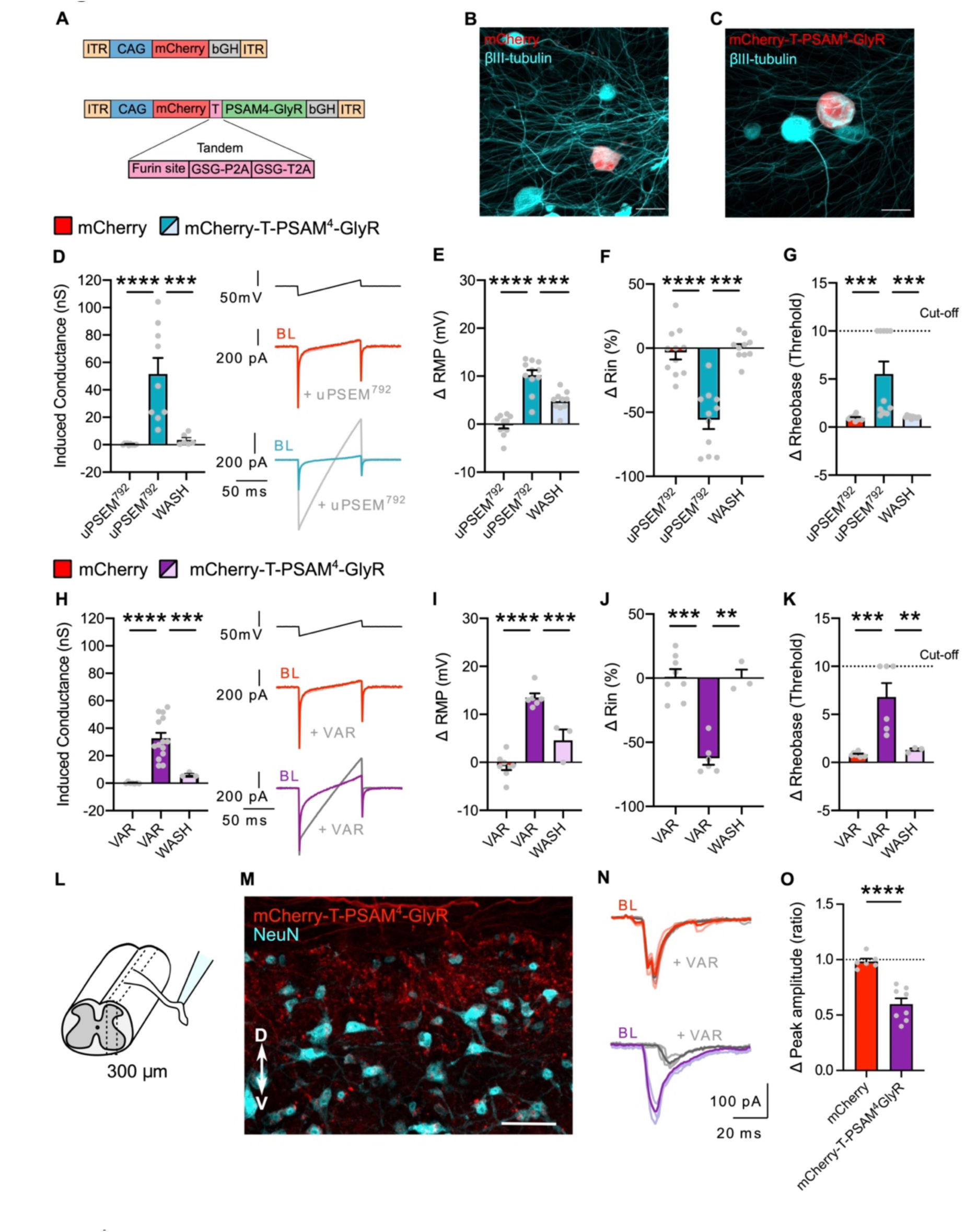
PSAM^4^-GlyR can reversibly silence mouse sensory neurons and reduce synaptic transmission. **(A)** Schematic representation of the constructs used in this study: mCherry and mCherry-T-PSAM^4^-GlyR. **(B-C)** Example images of dissociated sensory neurons transduced with mCherry **(B)** and mCherry-T-PSAM^4^-GlyR **(C)** after 2 days *in vitro* (2 DIV) (Scale bar = 50 μm). **(D)** uPSEM^792^ (10 nM) increased membrane conductance in PSAM^4^-GlyR expressing neurons. Induced conductance returned to baseline levels upon agonist washout. Inset shows representative current traces obtained by the application of voltage ramps used to determine membrane conductance. (mCherry: n = 10 cells; mCherry-T-PSAM^4^-GlyR: n = 9 cells). **(E-G)** Change in resting membrane potential (RMP) **(E)** input resistance **(F)** and rheobase **(G)** after uPSEM^792^ administration and washout. The cut-off rheobase value was defined as 10 times the threshold rheobase before uPSEM^792^ treatment. (mCherry: n = 11 cells; mCherry-T-PSAM^4^-GlyR: n = 11 cells). **(H)** Increased membrane conductance in PSAM^4^-GlyR neurons after varenicline (VAR; 20 nM) administration. Inset on the right shows representative traces of voltage ramp to determine membrane conductance. (mCherry: n = 7 cells; mCherry-T-PSAM^4^-GlyR: n = 14 cells). **(I-K)** Change in resting membrane potential (RMP) **(E)** input resistance **(F)** and rheobase **(G)** after varenicline administration. All changes induced by varenicline were reversible. (mCherry: n = 8 cells; mCherry-T-PSAM^4^-GlyR: n = 6 cells). D-G, One-way ANOVA with Tukey Post-hoc, ** P < 0.01, *** P < 0.001, **** P < 0.0001). **(L)** Schematic representation of the experimental procedure for spinal cord slices. **(M)** Example image of a parasagittal slice used for recording, showing mCherry-T-PSAM^4^-GlyR expressing terminals in the dorsal horn of the spinal cord. D: dorsal; V: ventral (Scale bar = 50 μm). **(N)** Representative postsynaptic currents evoked by dorsal root stimulation (ePSCs) recorded from superficial dorsal horn neurons obtained from mCherry-transduced animals (red) or mCherry-T-PSAM^4^-GlyR animals (purple) before and after application of varenicline (20 nM; grey). **(O)** Change of ePSC amplitude after application of varenicline (mCherry: n = 6 cells; mCherry-T-PSAM^4^-GlyR: n = 8 cells. Unpaired t-test, **** P<0.0001). All data are expressed as mean ± S.E.M.

### Activation of PSAM^4^-GlyR silences electrical activity of mouse sensory neurons

To assess the suitability of PSAM^4^-GlyR as a chemogenetic silencer in sensory neurons, we expressed mCherry-alone or mCherry-T-PSAM^4^-GlyR in dorsal root ganglion (DRG) sensory neurons acutely dissociated from mice (Fig. 1B-C), and performed florescence targeted whole-cell patch clamp recordings. We first determined the optimal dose of agonists (uPSEM^792^ or varenicline) to activate PSAM^4^-GlyR in *in vitro* experiments. Activating PSAM^4^-GlyR with the ultrapotent-designer agonist uPSEM^792^, produced a large increase in membrane conductance at very low doses. Administration of 2 nM uPSEM^792^ produced a small increase in membrane conductance which did not reach significance (8.9 ± 3.5 nS; p=0.2). However, progressively increasing uPSEM^792^ concentration significantly increased membrane conductance with 10 nM (38.7 ± 7.4 nS), 20 nM (43.4 ± 8.6 nS) and 100 nM (42.3 ± 8.3 nS) compared to mCherry control neurons (Fig. S2A). For all further *in vitro* experiments, we selected 10 nM uPSEM^792^ as appropriate for PSAM^4^-GlyR activation. As in cortical neurons (*31*), very low doses of varenicline were required to increase membrane conductance in PSAM^4^-GlyR-expressing cells. We found that administration of 2 nM varenicline did not produce a significant increase in membrane conductance (3.7 ± 1.5 pS) compared to control neurons expressing only mCherry, but increasing the dose to 10 nM increased the membrane conductance to 27.8 ± 4.8 nS which was significantly different from mCherry control neurons (p = 0.0005) (Fig. S2B). This increase was not significantly different with 20 nM (29.1 ± 4.1 nS; p < 0.0001) or 100 nM (24.2 ± 4.7 nS; p = 0.001) varenicline (Fig. S2B). We therefore decided to use 20 nM varenicline in all subsequent experiments to ensure sufficient PSAM^4^-GlyR activation.

We next addressed whether such an increase in membrane conductance impacts sensory neuron excitability. Sensory neurons were recorded in whole-cell current-clamp configuration, with a chloride load in the recording pipette (30 mM) to mimic the high intracellular chloride concentration maintained by sensory neurons (*35, 36*). Under these conditions, activation of PSAM^4^-GlyR with uPSEM^792^ (Fig. 1D) or varenicline (Fig. 1H) resulted in membrane potential (*V*_m_) depolarization (Fig. 1E and I), according to the chloride reversal potential estimated with the Nernst equation (*E_Cl_* = -41 mV). Depolarisation of *V*_m_ by agonist administration in PSAM^4^-GlyR neurons did not produce any AP firing, but was efficient in silencing sensory neurons via a large and significant decrease in input resistance (Fig. 1F and J), compared to mCherry control neurons. Furthermore, agonist administration increased the current required to generate an AP (rheobase) in PSAM^4^-GlyR expressing neurons, but not in control neurons (Fig.1 G and K). All the effects of agonist induced PSAM^4^-GlyR silencing were reversible *in vitro*; we observed washout and return to baseline of all parameters after 15 mins of agonist wash (Fig 1D-K). These results suggest that activation of PSAM^4^-GlyR potently, effectively and reversibly silences sensory neuron activity through large shunting chloride conductances.

### PSAM^4^-GlyR activation suppresses synaptic transmission to dorsal horn neurons

To determine whether silencing of neuronal activity observed at the soma translates into effective silencing at the presynaptic terminals in the spinal cord, we measured synaptic transmission in a parasagittal spinal cord slice preparation. To ease delivery of the construct *in vivo*, we generated AAV serotype 9 particles containing either mCherry-only or mCherry-T-PSAM^4^-GlyR. Spinal cord slices were obtained from virally-transduced mice, that had received neonatal subcutaneous AAV (which targets only the PNS (*37*)). Whole-cell patch clamp recordings were obtained from superficial dorsal horn neurons and monosynaptic excitatory post-synaptic currents evoked by stimulation of dorsal roots (eEPSCs) were measured (Fig. 1L-M). Administration of varenicline did not change the magnitude of eEPSCs in control neurons obtained from mCherry-transduced animals, whereas it greatly decreased the magnitude of postsynaptic responses obtained from neurons from PSAM^4^-GlyR expressing animals (Fig. 1N-O). This suggests that powerful silencing of electric activity at the soma and central terminals effectively interferes with the relay of information to the spinal cord.

### Activation of PSAM^4^-GlyR leads to decreased thermal and mechanical sensitivity

Once we determined the effective neuronal silencing capacity of PSAM^4^-GlyR activation *in vitro* and *ex vivo*, we then explored whether PSAM^4^-GlyR activation silences sensory behaviours *in vivo*. We virally delivered PSAM^4^-GlyR by intrathecal (i.t.) injection to selectively transduce sensory neurons, while avoiding transduction of neurons in the spinal cord (*38*). After 6-8 weeks post i.t. injection we measured the effect of PSAM^4^-GlyR activation upon thermal sensitivity using the Hargreaves radiant heat source (Fig. S3A). We established a time course after an intraperitoneal (i.p.) injection of uPSEM^792^ (1mg/kg), where we saw a decrease in thermal sensitivity at 2 hours from treatment which returned to baseline after 5 hours. One week after uPSEM^792^ administration, thermal thresholds were comparable to baseline levels which enabled reactivation of PSAM^4^-GlyR by subsequent administrations (Fig. S3B). Treatment with a higher dose of uPSEM^792^ (5mg/kg) produced a stronger decrease in thermal sensitivity after 1 hour, again peaking at 2 hours from i.p. injection, which recovered after 5 hours (Fig. S3C-D). From these data we were confident that our silencing system was functional *in vivo*.

From a clinical translation perspective, the significance of gene therapy relies on long-term expression of the transgene. To look at the stability of PSAM^4^-GlyR expression over time, we tested again the same behavioural cohort 9-10 months after viral delivery. Following uPSEM^792^ administration, PSAM^4^-GlyR expressing mice became hyposensitive to punctate mechanical stimuli (von Frey; Fig.2B), noxious mechanosensation (pinprick; Fig. 2C.) and light touch (brush; Fig. 2D). In addition, mice also became hyposensitive to noxious hot and cold thermal stimuli (Hargreaves; Fig. 2E. and dry ice; Fig. 2F) after treatment with uPSEM^792^. Control mice were unaffected by uPSEM^792^ administration in all assays. Complete washout of uPSEM^792^ was achieved and all changes in sensory sensibility returned to baseline (Fig. 2B-F). This allowed reactivation of PSAM^4^-GlyR with varenicline (0.3mg/kg, the more clinically relevant PSAM^4^-GlyR agonist), which produced a similar behavioural hyposensitivity to sensory stimuli throughout all assays in PSAM^4^-GlyR expressing mice, which was completely reversible (Fig 2G-K); the same dose of varenicline had no effects on the sensory function of control mice (Fig. 2G-K). In contrast, varenicline administration produced no proprioceptive changes in PSAM^4^-GlyR expressing mice, as they behaved similar to control mice during the beam task (Fig. S4A-D). Taken together, PSAM^4^-GlyR expression and function remained stable in the long-term and yielded powerful and reversible silencing of sensory behaviours upon activation with either the designer or the clinically-viable PSAM^4^-GlyR agonist.

**Fig. 2.**
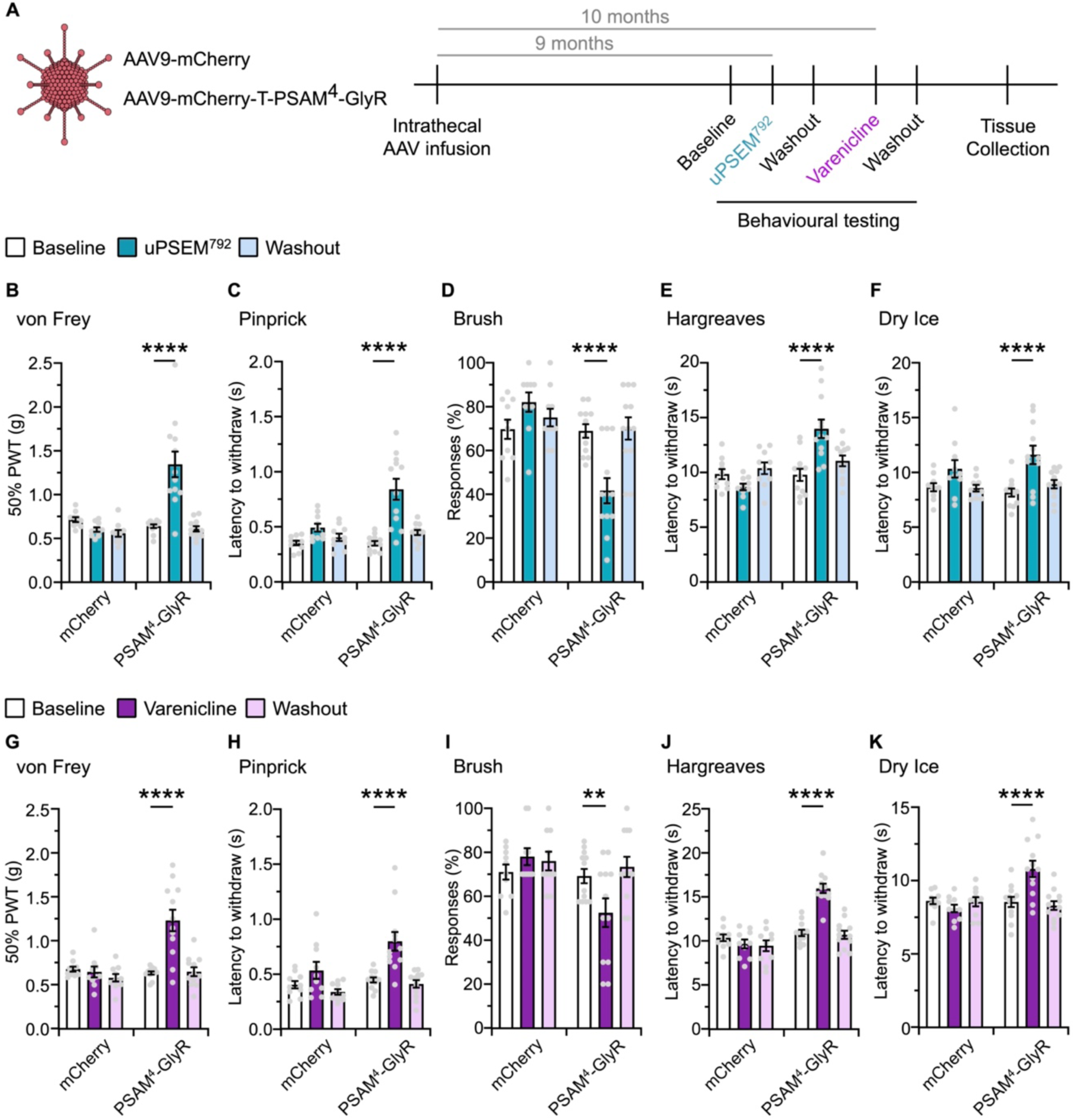
Robust, reversible, and repeatable silencing of acute sensory behaviours via agonist-induced activation of PSAM^4^-GlyR. (**A**) Timeline of the experimental design. Intrathecal infusion of AAV-mCherry or mCherry-T-PSAM^4^-GlyR followed by baseline sensory testing, and re-testing during or post uPSEM^792^ or varenicline. (**B – D**) Mechanical sensory testing was unchanged in mCherry expressing mice, but PSAM^4^-GlyR expressing mice were significantly hypo-sensitive to von Frey (**B**), Pinprick (**C**) and Brush (**D**) stimuli following uPSEM^792^ which was reversible. (**E** and **F**) Thermal sensory testing was normal in mCherry expressing mice, but PSAM^4^-GlyR expressing mice were significantly hypo-sensitive to Hargreaves (**E**) and Dry Ice (**F**), following uPSEM^792^. All uPSEM^792^ induced silencing was reversible. (**G-I**) Mechanical sensory testing was normal in mCherry expressing mice, but PSAM^4^-GlyR expressing mice were significantly hypo-sensitive to von Frey (**G**), Pinprick (**H**) and Brush (**I**) stimuli following the clinical agonist varenicline. (**J** and **K**) Thermal sensory testing was unchanged in mCherry expressing mice, but PSAM^4^-GlyR expressing mice were significantly hypo-sensitive to Hargreaves (**J**) and Dry Ice (**K**) following varenicline. All varenicline induced silencing was reversible. (mCherry: n = 10 mice, PSAM^4^-GlyR: n = 12 mice, all data sets RM-two way ANOVA, post-hoc Bonferroni test, ** P < 0.01, **** P < 0.0001). Data expressed as mean ± S.E.M.

### Stable expression of PSAM^4^-GlyR in DRG sensory neurons

Expression of PSAM^4^-GlyR was verified after behavioural experiments to confirm long-term expression of the construct after i.t. viral delivery. We found that our approach transduced different sensory neurons in an unbiased manner (Fig. 3A). Approximately 50% of neurons in L4 DRG displayed a clear fluorescent signal, without antibody amplification (Fig. 3B). AAV-mCherry-T-PSAM^4^-GlyR co-expressed with specific subpopulation markers (Fig. 3C-H). Expression was comparable across different subpopulations, but with lower expression in the TH subpopulation (Fig. 3H). We also found extensive distribution of PSAM^4^-GlyR terminals in superficial and deep layers of the spinal cord, but no spinal cord neurons were observed to express mCherry-T-PSAM^4^-GlyR (Fig. 3I). Importantly, we also found no increase of the injury marker ATF3, in DRG neurons, indicating that long-term expression of PSAM^4^-GlyR was not associated with overt neuronal damage (Fig. S5A-B). Our results suggest that i.t. viral delivery of PSAM^4^-GlyR selectively transduced different types of DRG neurons with comparable efficiency, and remained stable after many months.

**Fig. 3.**
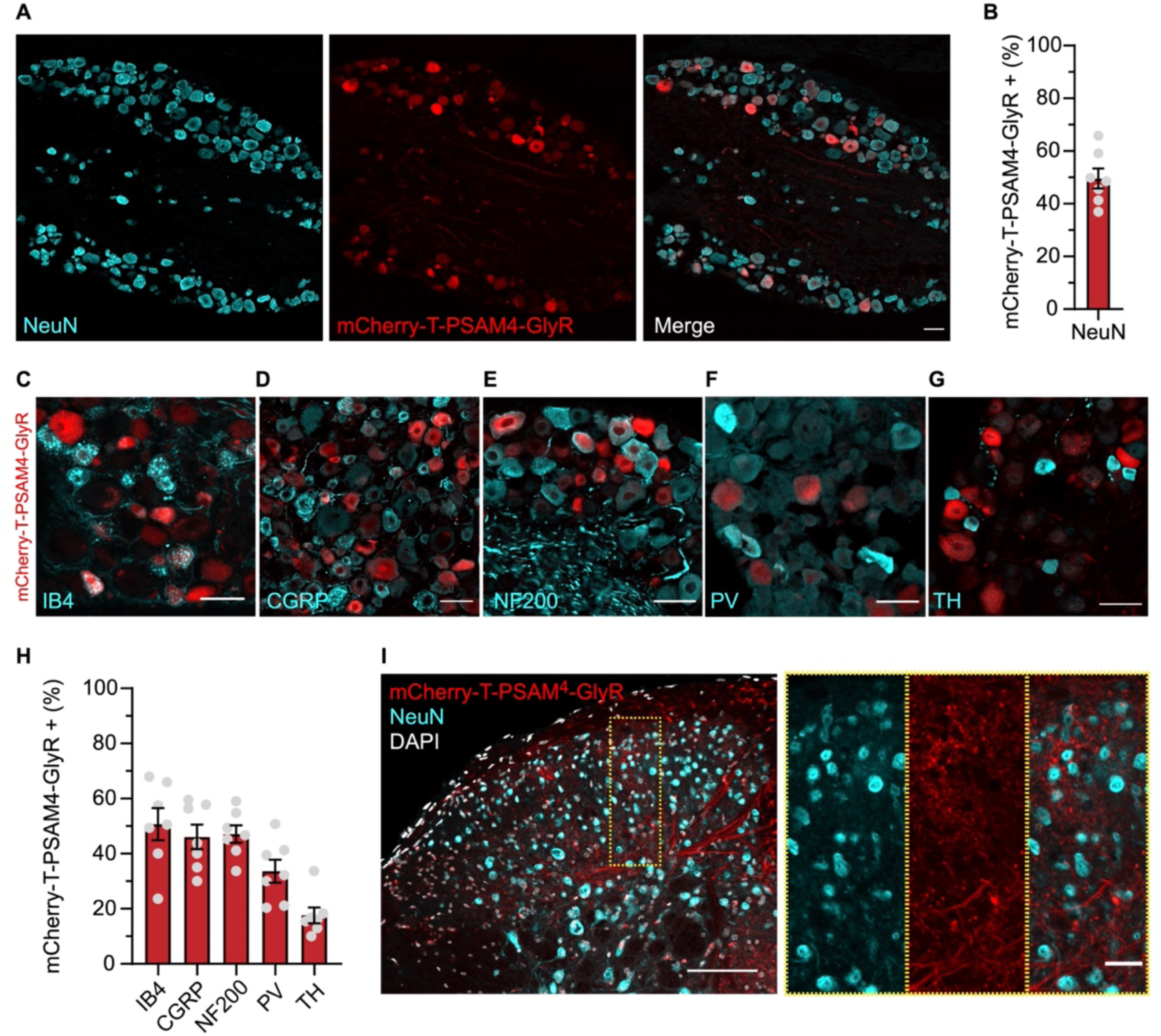
Long-term, stable expression of AAV-mCherry-T-PSAM^4^-GlyR selectively in sensory neurons. (**A**) Example image of DRG neurons transduced by AAV-mCherry-T-PSAM^4^-GlyR 11-months post i.t. injection (Scale bar 50 um) (**B**) Quantification of NeuN positive neurons that are also mCherry-T-PSAM^4^-GlyR positive (n = 7 mice, 1631/3229 neurons) (**C – G**) Example images of DRG neuron subpopulation markers, IB4 (**C**), CGRP (**D**), NF200 (**E**), PV (**F**), TH (**G**), and their co-expression with mCherry-T-PSAM^4^-GlyR (Scale bars 25 um). (**H**) Percentage of each DRG neuron subpopulation that co-express mCherry-T-PSAM^4^-GlyR (n = 7 mice, (IB4: 480/973 neurons, CGRP: 435/967 neurons, NF200: 479/1014 neurons, PV: 69/219 neurons, TH: 51/261 neurons). (**I**) Example image of mCherry-T-PSAM^4^-GlyR positive afferents entering and terminating in the dorsal horn of the spinal cord (Scale Bar 100 µm, insert scale bar 25 µm). Data mean ± S.E.M.

### PSAM^4^-GlyR mediated silencing suppressed pain-related behaviours in inflammatory pain models

Following inflammation or injury it is common for nociceptors to become sensitized, showing hyperexcitability, which is key in driving pain (*3*). Therefore, we next asked if our humanised silencing system could be applied to pathological pain conditions. We first chose a chemical, inflammatory-like pain model where formalin was injected into the hindpaw of mice expressing either AAV-mCherry or AAV-mCherry-T-PSAM^4^-GlyR. All mice were dosed with varenicline 1 hr prior to the formalin assay and assessment of nocifensive behaviours were conducted. We found that mice expressing PSAM^4^-GlyR exhibited significantly less pain-related behaviours in the 1^st^ phase of the formalin assay, compared to control mice (Fig. S6A-B). These results were encouraging so we undertook a more clinically relevant model of inflammatory joint pain (Fig. 4A). In this cohort of mice, we selectively targeted knee-innervating afferents using intra-articular (i.a.) AAV serotype PHP.S, as reported previously (*24*) (Fig. 4B). Mice received either AAVPHP.S-eGFP or AAVPHP.S-mCherry-T-PSAM^4^-GlyR and 4 weeks later all mice received an i.a. injection of the sensitising agent Complete Freund’s Adjuvant (CFA). As expected, all mice that received CFA developed knee swelling, ipsilateral to the injection (Fig. 4C). Mice were tested prior to, 24 hrs post CFA, and 1 hr following varenicline induced silencing. Consistent with previous findings (*24*), across all testing points motor function remained intact and no deficit in rotarod performance was observed (Fig. 4D). Following CFA, in both eGFP and PSAM^4^-GlyR groups, the mechanical pressure threshold of the contralateral knee was unchanged (Fig. 4E), but the ipsilateral knee became significantly hypersensitive (Fig. 4F). Varenicline was then given to see if PSAM^4^-GlyR mediated silencing could recover inflammatory-induced mechanical hypersensitivity. Whereas no change in mechanical pressure hypersensivity occurred in the control group, a significant partial recovery towards baseline threshold was observed in PSAM^4^-GlyR expressing mice (Fig. 4F). Inflammatory joint pain can alter naturalistic-like behaviours in mice that can be used as a readout of spontaneous pain (*5, 24*). Following i.a. CFA all mice dug fewer burrows and spent less time digging, suggesting they were in a pain-like state (Fig. 4G-H). Following varenicline, control mice continued to exhibit reduced digging, whereas PSAM^4^-GlyR expressing mice exhibited increased digging (Fig. 4G-H). Collectively, our data indicate that PSAM^4^-GlyR mediated silencing of knee innervating afferents can reduce both evoked and non-evoked pain-related behaviours associated with knee inflammation.

**Fig. 4.**
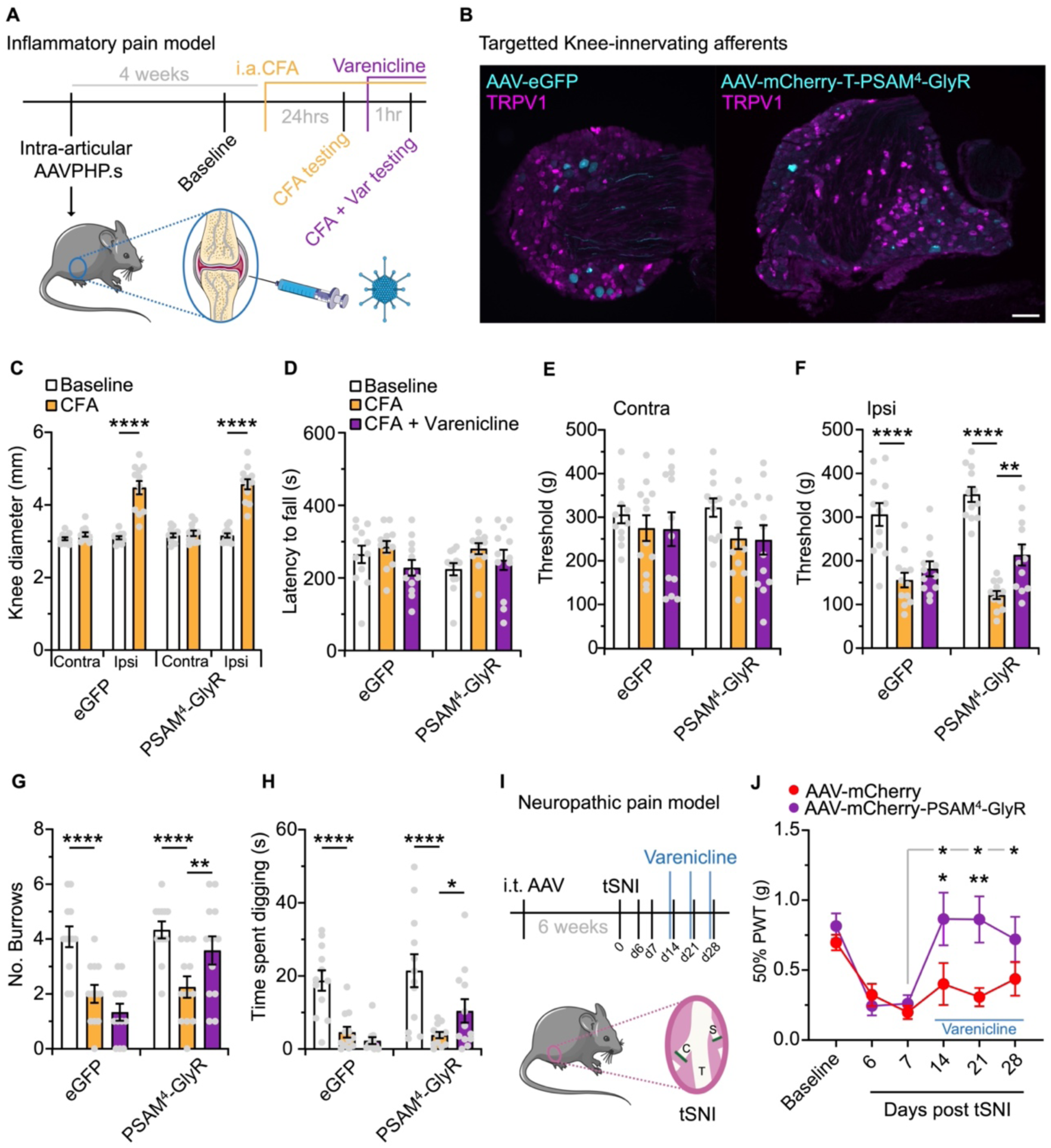
PSAM^4^-GlyR mediated silencing of inflammatory-joint and neuropathic pain. (**A**) Schematic of the experimental timeline of targeting joint afferents with AAVPHP.S, baseline testing, i.a. CFA and behavioural test pre- and post-varenicline. (**B**) Example images of L4 DRG neurons transduced with either AAVPHP.S-eGFP or AAVPHP.S-mCherry-T-PSAM^4^-GlyR following i.a. injection (scale bar = 100 µm). (**C**) i.a. CFA injection resulted in significant knee swelling only in ipsilateral joints (eGFP: n = 12 mice, PSAM^4^-GlyR = 12 mice, RM two-way ANOVA, post hoc Bonferroni test, **** P<0.0001). (**D**) Motor function was not altered by CFA or varenicline induced silencing. (**E**) The mechanical pressure threshold of contralateral knee joints were normal throughout the experiment. (**F**) Ipsilateral knees in both groups become significantly hypersensitivity to mechanical pressure following CFA. Only in the PSAM^4^-GlyR group was mechanical hypersensitivity partially reversed following varenicline. (**G** and **H**) Following CFA, mice from both groups burrowed less (**G**) and spent less time digging (**H**). Varenicline recovered the number of burrows and time spent digging in PSAM^4^-GlyR expressing mice. (D-H, eGFP: n = 12 mice, PSAM^4^-GlyR = 12 mice, RM two-way ANOVA, post hoc Bonferroni tests compared to CFA ,* P < 0.05, ** P < 0.01, **** P<0.0001). (**I**) Depiction of the neuropathic pain model experimental timeline. Intrathecal targeting of sensory neurons, followed by tSNI and mechanical testing pre- and post-varenicline. C: common peroneal; T: tibial; S: sural. (**J**) Mice from both groups became hypersensitive to mechanical stimuli following injury. Mice expressing PSAM^4^-GlyR recovered from mechanical hypersensitivity following varenicline compared to day 7 (grey line) and compared to mCherry expressing mice (mCherry: n = 10 mice, PSAM^4^-GlyR = 12 mice, RM two-way ANOVA, post hoc Bonferroni tests compared to D7 (grey line) or between groups, * P < 0.05, ** P < 0.01). Data expressed as mean ± S.E.M

### Activation of PSAM^4^-GlyR suppressed mechanical hypersensitivity after nerve injury

Injury, disease or genetic mutations affecting the nervous system can result in neuropathic pain. Injury to the nervous system also entails significant sensory neuron hyperexcitability, which is thought to drive neuropathic pain (*6–8*). Thus, we next asked, if PSAM^4^-GlyR-mediated silencing of sensory afferents can treat mechanical allodynia associated with neuropathic pain. We chose the tibial spared nerve injury model (tSNI) and measured the mechanical withdrawal thresholds of mice (*39*), that had previously received i.t. AAV-mCherry or AAV-mCherry-T-PSAM^4^-GlyR (Fig. 4I). All mice became hypersensitive to mechanical stimuli by day 7 (Fig. 4J). On days 14, 21 and 28 after tSNI, we treated all mice with the clinically relevant PSAM^4^-GlyR agonist varenicline, 1 hr before mechanical testing. We found that mechanical hypersensitivity recovered following varenicline treatment in mice transduced with AAV-mCherry-T-PSAM^4^-GlyR, but not in control mice (Fig. 4J). Therefore, PSAM^4^-GlyR activation effectively silences pain-related behaviour after nerve injury.

### Activation of PSAM^4^-GlyR silences excitability in human-derived sensory neurons

So far, we have shown the suitability of PSAM^4^-GlyR as an effective chemogenetic silencer in animal models. To demonstrate its applicability as a fully humanised system, we virally expressed PSAM^4^-GlyR in human induced pluripotent stem-cell-derived sensory neurons (hiPSC-SNs). After differentiation and maturation, we transduced hiPSC-SNs with AAV9-mCherry or AAV9-mCherry-T-PSAM^4^-GlyR (Fig. 5A). At least four weeks after viral delivery, transduced cells exhibited very bright fluorescence which we used for targeted whole-cell patch clamp recordings (Fig. 5B). We found a robust increase in membrane conductance after administration of both uPSEM^792^ and varenicline (Fig. 5C and G). We also examined the effect of activating PSAM^4^-GlyR on neuronal excitability. Both uPSEM^792^ and varenicline produced a significant depolarization of the resting membrane potential (Fig. 5D and H), similar to what we observed in mouse sensory neurons. Both PSAM^4^-GlyR agonists lead to a decrease in input resistance (Fig. 5E and I), which limited the capacity of hiPSC-SNs to generate APs. Indeed, most neurons reached the threshold cut-off for rheobase (Fig. 5F and J). We also measured the firing patterns of hIPSC-SNs following sustained current injections. Many cells exhibited repetitive firing in response to progressive depolarizing currents, which was abolished upon PSAM^4^-GlyR mediated silencing (Fig. S7A-B). These results suggest that PSAM^4^-GlyR can produce powerful silencing of neuronal activity not only in mice, but also human sensory neuron models, validating its candidacy as a gene therapeutic.

**Fig. 5.**
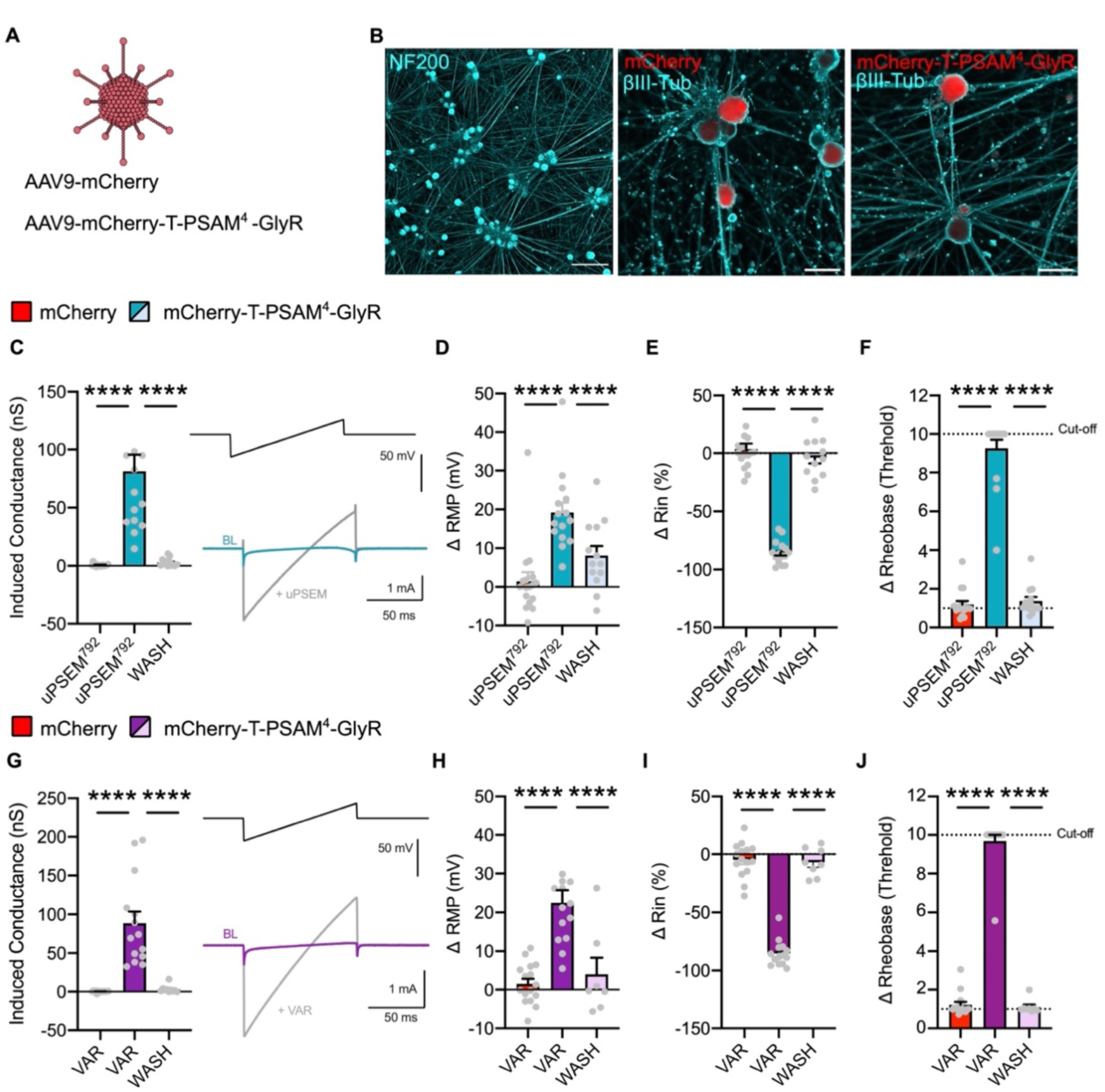
PSAM^4^-GlyR mediated silencing of human iPSC derived sensory neurons. **(A)** AAV9-mCherry or AAV9-mCherry-T-PSAM^4^-GlyR were used to virally transduce sensory neurons derived from human induced pluripotent stem cells (hiPSC-SNs). **(B)** Example images showing mature hiPSC-SNs (*left*) and mCherry labeling in AAV-mCherry (*middle*) and AAV-mCherry-T-PSAM^4^-GlyR (*right*) transduced neurons. **(C)** Robust change in membrane conductance after uPSEM^792^ application. Inset shows representative traces of voltage clamp recordings used to measure membrane conductance. **(D-F)** Change in membrane potential **(B)** input resistance **(E)** and rheobase **(F)** after uPSEM^792^ administration. All changes were reversible upon agonist washout (mCherry: n = 16 cells; mCherry-T-PSAM^4^-GlyR: n = 15 cells. One-way ANOVA with Tukey post-hoc, **** P<0.0001). **(G)** Varenicline application produced a change in membrane conductance. Insets show representative currents of the change in membrane conductance after agonist administration. **(H-J)** Changes in membrane potential **(H)** input resistance **(I)** and rheobase **(J)** after varenicline treatment. All changes were reversible upon varenicline washout (mCherry: n = 15 cells; mCherry-T-PSAM^4^-GlyR: n = 14 cells. One-way ANOVA with Tukey post-hoc, **** P<0.0001). Rheobase cut-off was defined as 10 times the baseline rheobase threshold. All data are expressed as mean ± S.E.M.

### Silencing spontaneous activity in a clinical model of neuropathic pain

Spontaneous activity (SA) in sensory neurons is thought to be one of the key pathophysiological drivers of peripheral neuropathic pain conditions, and is often well correlated with spontaneous pain experienced by patients (*40, 41*). Gain-of-function (GoF) mutations in *SCN9A*, the gene that encodes the voltage-gated sodium channel subunit Na_V_1.7, are associated with inherited erythromelalgia (IEM, presenting with pain and erythema of the extremities) and increased sensory neuron spontaneous activity *in vivo* and *in vitro* (*42*). To test the efficacy of our humanised PSAM^4^-GlyR channel in a human neuropathic pain model, we generated hiPSC-SN from iPSCs originating from a patient with IEM (GoF mutation in Na_V_1.7-F1449V), previously reported (*43*) (Fig. 6A). We treated mature control and IEM hiPSC-SNs with our viruses (AAV-mCherry or AAV-mCherry-T-PSAM^4^-GlyR) and four weeks later measured SA in a cell-attached patch clamp configuration to avoid dialysing the internal milieu of the cells, at 32°C (Fig. 6A). We found low levels of SA in cells differentiated from control patients (0.3 ± 0.2 Hz; Fig. 6B), whereas cells differentiated from the IEM patient showed higher SA (6.5 ± 3.7 Hz; Fig. 6C), similar to what has been previously reported (*43, 44*). We conducted a population-based analysis of hiPSC-SNs from healthy controls and IEM that were transduced with mCherry or mCherry-T-PSAM4-GlyR to determine whether PSAM^4^-GlyR activation by varenicline could suppress the SA observed in the IEM line. Healthy control mCherry expressing neurons exhibited a low incidence of SA (14.29%; Fig. 6D) compared to the high incidence of SA in IEM neurons (30.3%; Fig. 6E). Varenicline treatment decreased the frequency of firing activity and the proportion of spontaneously active neurons in both control (0.04 ± 0.05 Hz; 3.33%) and IEM neurons (0.004 ±0.02 Hz; 14.71%) that expressed PSAM^4^-GlyR (Fig. 6F and G). Importantly, varenicline treatment decreased the proportion of IEM neurons with SA in PSAM^4^-GlyR -expressing neurons, closer to the proportion observed in mCherry expressing control neurons (Fig. 6D). Our results highlight the translational potential of PSAM^4^-GlyR in suppressing ectopic activity in human sensory neurons, a key driver of neuropathic pain.

**Fig. 6.**
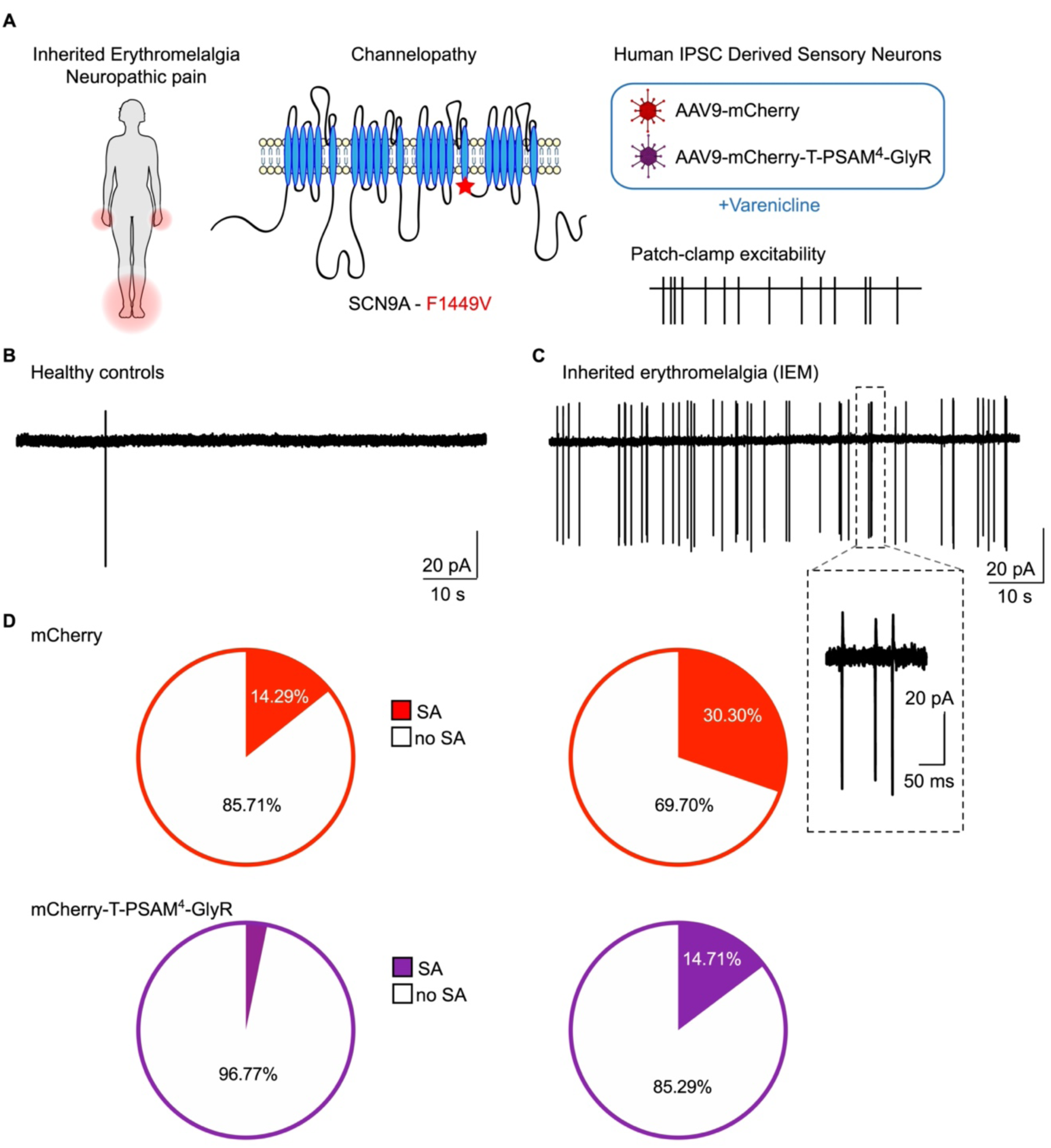
Silencing of spontaneous activity in a human neuropathic pain model. **(A)** Schematic representation the experimental design. Patients with inherited erythromelalgia (IEM) exhibit pain and erythema in the extremities. The schematic representation of the Nav1.7 channel α-subunit shows the location of the point mutation (F1449V), used in this study. Transduction of hiPSC-SN with AAV-mCherry and AAV-mCherry-T-PSAM^4^-GlyR to study the effect of silencing on spontaneous activity in control and IEM patients. **(B)** Spontaneous activity recorded in cell-attached configuration in hiPSC-SNs derived from a healthy control patient. **(C)** Cell-attached recording showing spontaneous activity obtained in hiPSC-SNs derived from a patient with IEM. Inset shows an amplification of a region showing bursting events. **(D)** Proportion of AAV-mCherry transduced neurons obtained from healthy control patients showing no spontaneous activity (no SA; n = 24, white) and spontaneous activity (SA; n = 4, red). **(E)** Proportion of AAV-mCherry transduced neurons obtained IEM patients showing no spontaneous activity (no SA; n = 23, white) and spontaneous activity (SA; n = 10, red). χ^2^ = 6.9, P = 0.009 **(F)** Proportion of AAV-mCherry-T-PSAM^4^-GlyR transduced neurons obtained from healthy control patients showing no spontaneous activity (no SA; n = 30, white) and spontaneous activity (SA; n = 1, purple). **(G)** Proportion of AAV-mCherry-T-PSAM^4^-GlyR transduced neurons obtained from IEM patients showing no spontaneous activity (no SA; n = 29, white) and spontaneous activity (SA; n = 5, purple). χ^2^ = 3.9, P = 0.04.

## DISCUSSION

In this study, we show that expression of the human chemogenetic system PSAM^4^-GlyR in mice can silence the activity of sensory neurons, and reduce behavioural hypersensitivity in inflammatory and neuropathic pain models. Viral delivery results in long-term stable expression of PSAM^4^-GlyR in different populations of sensory neurons, and can be repeatedly activated *in vivo*. We also used PSAM^4^-GlyR to silence the excitability of human sensory neurons and to decrease ectopic discharge in patient-derived sensory neurons with a GoF Na_V_1.7 mutation causing IEM. We propose that PSAM^4^-GlyR is a promising humanised gene therapy that can be used to silence ectopic activity and hyperexcitability associated with multiple chronic pain disorders, such as inflammatory joint pain and neuropathic pain.

The inhibitory effect of opening chloride channels is highly dependent on the chloride gradient across the membrane (*45*). Unlike most neurons, sensory neurons maintain a high intracellular chloride concentration, that leads to chloride efflux upon chloride channel activation. However, the ensuing membrane depolarization does not produce neuronal firing of sensory neurons. Instead, the PSAM^4^-GlyR mediated chloride currents in these neurons produce strong inhibition of neuronal activity. This is likely a cumulative mechanism of two key factors. Firstly, PSAM^4^-GlyR activation drives the membrane potential toward *E*_Cl,_ which would inactivate voltage-gated sodium channels (*46–49*). Secondly, we see a dramatic reduction in membrane resistance following PSAM^4^-GlyR activation. Together, we believe these factors render sensory neurons unable to generate and propagate APs due to chloride mediated shunting inhibition (*50*). Unlike sensory neurons, CNS neurons are very susceptible to the smallest changes in intracellular chloride, which affect the very nature of inhibition (hyperpolarising/depolarising), and may lead to the collapse of the chloride gradient, especially upon repetitive activation (*51, 52*). Thus, using chloride conductances as a means to suppress neuronal activity, may be particularly suited and effective in sensory neurons, compared to other neurons (*32, 38*). Interestingly, enhanced chloride loading in sensory neurons may also occur after inflammation and nerve injury, which depolarises *E*_Cl_ further, leading to spiking (*49*), especially in combination with hyperexcitability (*53*). Yet, even in these pathological conditions, we believe any depolarisation due to sodium ion entry can be counterbalanced by chloride flux, thus maintaining the shunting inhibition (*50*). Indeed, we found that activation of PSAM^4^-GlyR effectively and reversibly silences activity in sensory neurons, and silences pain behaviour even in pathological conditions such as inflammation and nerve injury.

The idea of using PSAM^4^-GlyR for therapeutic treatment of hypersensitivity in neuropathic pain conditions is of great interest. We showed that expressing the chemogenetic channel in hiPSC-SNs led to strong inhibition and complete silencing, the magnitude of which was comparable to the silencing observed in rodent neurons. Yet, this is the first report of using chemogenetics as a gene therapy to silence the activity of human sensory neurons from a cellular model of hyperexcitability and neuropathic pain. Deriving hiPSC-SNs from healthy and people living with disease is a powerful and accessible model system to test therapeutic design (*54*). We used this system to model inherited erythromelalgia, an inherited channelopathy, which in people leads to intense episodes of spontaneous, burning pain, which is often triggered by warming (*42*). We derived hiPSC-SNs from a patient with the F1449V GoF mutation in *SCN9A*. This mutation has been shown to result in GoF with a hyperpolarizing shift in the voltage dependence of activation of Na_V_1.7 and a depolarizing shift in steady-state inactivation resulting in lowered threshold for single APs and high frequency firing when expressed in rodent DRG neurons (*55*). We tested the efficacy of varenicline activation of PSAM^4^-GlyR in silencing the ectopic activity seen in hiPSC-SNs. Our data demonstrate that we can silence the aberrant activity associated with this neuropathic pain condition emphasizing the therapeutic potential of PSAM^4^-GlyR.

Previous efforts to decrease hyperexcitability in hiPSC-SNs derived from IEM patients have focused on blocking Na_V_1.7 channels or countering their activity with dynamic clamp, both of which have produced encouraging results (*43, 56*). Translating these treatments to humans should provide pain amelioration without side effects, because Nav1.7 is preferentially expressed in the peripheral system, and is crucial for sensory neuron activity (*42*). However, targeting a single molecular target has not fared well in the clinic (*19*), probably because of degeneracy – changes in many ion channels and molecules may produce hyperexcitability (*57*) and attribution of hyper-excitability and clinical pain to a single channel mutation is rare. Our strategy of using PSAM^4^-GlyR to silence sensory neurons by-passes this problem, by efficiently suppressing ectopic activity irrespective of the underlying molecular mechanisms.

Important concerns have emerged regarding the clinical application of current chemogenetic systems. In the case of designer receptors exclusively activated by designer drugs (DREADDs), these are associated with their specific agonist CNO which is rapidly converted into neuroactive compounds that can act on other endogenous targets (*28, 58*), as well as producing ligand-independent changes in ion channels and second-messenger signaling in sensory neurons (*26*). An alternative promising candidate is the engineered glutamate-gated chloride channel (GluCl) which has been shown to be effective in silencing sensory neurons (*38*). However, it was originated from an invertebrate protein, which may raise concerns about how well it would be tolerated and integrated in humans, especially with regard to immunogenicity (*38, 59*). Using PSAM^4^-GlyR may circumvent these concerns, the agonist varenicline has been used safely in humans as a treatment to aid smoking cessation (*60, 61*). This has the advantage of already being an FDA-approved drug for human use. Furthermore, the doses required to activate the channel are in the nanomolar range, well below those used in the clinic, a feature that arises due to the mutations made in the ɑ7 nicotinic receptor part of PSAM^4^-GlyR (*31*). Most importantly, PSAM^4^-GlyR is engineered from human proteins, the modified human ACh nicotinic and glycine receptors. While there is always concern when expressing non-endogenous proteins, regarding long-term safety and efficacy (*62, 63*), PSAM^4^-GlyR would likely be better tolerated compared to proteins originating from non-mammalian systems. Importantly, we demonstrated that a single i.t. injection to deliver AAV-mCherry-T-PSAM^4^-GlyR *in vivo* remained stable after 11 months without any evidence of neuronal damage.

Despite these promising features, there are some limitations to our present study. Therapeutic application of PSAM^4^-GlyR would require gene delivery into candidate patients. We have used i.t. and i.a. injection to deliver AAVs in animals *in vivo*. Viral vectors such as AAVs are widely used to deliver therapeutic genes (*64*). They are well tolerated in humans, and several therapeutic genes have been approved to use clinically (*65, 66*). However, delivering AAV-PSAM^4^-GlyR such that it effectively transduces sensory neurons *in vivo* in humans, remains a major challenge; i.t. injection for AAV delivery is not a trivial procedure, but has been used for gene therapy (*67*); i.a. injection would be similar to procedures patients undergo as part of normal therapeutic treatment for arthritis conditions, such as corticosteroid injections. Large strides are being made in the design and delivery of novel AAV vectors with the intention of human gene therapies (*68–70*). Our rodent data support the use of AAVPHP.S encoding PSAM^4^-GlyR as a means of targeted delivery of our chemogenetic silencer directly to sensory neurons innervating the sensory target, i.e. the knee. This is of clinical interest as it is far less invasive than i.t. or intraneural injections and will result in selective silencing of sensory neurons driving the chronic pain state (*68*). The AAV-PSAM^4^-GlyR approach has shown broad utility across a number of diverse pain models with no observed long-term toxicity or motor dysfunction.

Further advances in focusing such a chemogenetic approach will likely arise from future targeting expression to specific sensory-neuron sub-populations. Indeed, we have good evidence that both low-threshold A-fibres, as well as C-fibre nociceptors are important in neuropathic pain (*71, 72*). However, we have yet to understand which sensory neuron subpopulations develop hyperexcitability and ectopic activity and contribute to specific features of inflammatory and neuropathic pain. Our viral strategy, using a generic promoter, led to broad expression in all sensory neuron subtypes, resulting in reduced pain, but also general hyposensitivity. We limited the expression of PSAM^4^-GlyR to L3-L5 DRG by i.t. injection, where input from the hindpaw is encoded and where we tested. Moving forward, generation of viral constructs dependent on Cre recombination, along with intersectional approaches, to effectively target specific sensory neuron subpopulations would be of significant benefit (*73, 74*). Alternatively, AAV capsid engineering or the use of human promoters specific to sensory neuron sub-populations could also be used to focus and restrict chemogenetic receptor expression (*75–77*).

Taken together, our study highlights the efficacy of PSAM^4^-GlyR in silencing sensory neuron hyperexcitability and demonstrates the translational potential of an effective, stable and reversible human-based chemogenetic system for the treatment of pain.

## MATERIALS AND METHODS

### Animals

All mice were group-housed in individually ventilated cages with free access to food and water, in humidity and temperature controlled rooms with a 12 hr light-dark cycle, in a pathogen free facility. All animal procedures adhered to the UK Home Office (Scientific Procedures) Act (1986) Amendment Regulations 2012, and were performed under a UK Home Office Project Licence. All animal experiments were carried out in accordance with University of Oxford Policy on the Use of Animals in Scientific Research. For experiments performed in Cambridge (knee joint inflammation and behavioural assessment) the University of Cambridge Animal Welfare and Ethical Review Body also approved all animal experiments.

The work within this study also conforms to the ARRIVE guidelines (*77*). All behavioural experiments were carried out by a blind experimenter on adult male and female mice. C57BL/6J wild type mice were sourced from biomedical services breeding unit at the University of Oxford and Envigo.

### Molecular cloning

pCAG PSAM4 GlyR IRES EGFP was a gift from Scott Sternson (Addgene plasmid # 119739) and was subcloned into a pAAV vector between AAV2 ITR sites. For our purpose this construct was too large for viral production. To shorten the construct and swap the reporter, we generated a plasmid iteration that replaced EGFP with mCherry, the IRES sequence with a Tandem sequence (Furin-GSG-P2A-GSG-T2A)(*78*). Our initial observations (data not shown) noted that the PSAM4-GlyR C-terminal domain is likely sensitive to reporter tagging/linking. Therefore, we opted to swap PSAM4-GlyR and mCherry ORFs to generate our final construct; pAAV-CAG-mCherry-Tandem-PSAM4-GlyR (now available on Addgene) In brief, to generate our insert, (mCherry-Tandem-PSAM4-GlyR), four fragments were generated (supplementary table X), digested using restriction enzymes (NEB) and ligated using Quick ligase (NEB); After fragment ligation, the product (2.2kb), was PCR amplified and inserted into a pAAV-CAG backbone using NheI and AgeI restriction sites. From this construct, we next generated our pAAV-CAG-mCherry control plasmid. We PCR amplified NheI-Kozak-mCherry, and inserted it into our pAAV-CAG backbone using NheI and AgeI restriction sites. All constructs were screened and sequence confirmed using Sanger sequencing.

| No. | Name | 5' restriction site | 3' restriction site | Source |
| --- | --- | --- | --- | --- |
| 1 | Kozak-(partial)mCherry | NheI | SbfI | Twist Bioscience |
| 2 | (Partial)mCherry | SbfI | SgrAI | Twist Bioscience |
| 3 | Tandem-(partial)PSAM4 | SgrAI | SexAI | Twist Bioscience |
| 4 | (Partial)PSAM4-GlyR | SexAI | AgeI | PCR from #119739* |

\* AgeI was introduced using a reverse primer overhang

### Viral production

pAAV-CAG-Kozak-mCherry-Tandem-PSAM4-GlyR and pAAV-CAG-Kozak-mCherry constructs were commercially packaged and serotyped with AAV9 capsid protein by the Viral Vector Facility (VVF), Neuroscience Centre Zurich (ZNZ), University of Zurich and ETH Zurich. Final titres for AAV9-CAG-mCherry-T-PSAM4-GlyR and AAV9-CAG-mCherry were 1.2 x 10^13^ vg/ml (vector genomes/milliliter) and 3.1 x 10^13^ vg/ml, respectively. pAAV-CAG-Kozak-mCherry-Tandem-PSAM4-GlyR was also commercially packaged into AAV serotype PHP.S at a final titre of 8.0 x 10^12^ vg/ml (VVF). AAVPHP.S-CAG-eGFP (v24-PHP.S) was purchased from VVF at a tire of 1.3 x 10^13^ vg/ml, and diluted to a final working tire of 8.0 x 10^12^ vg/ml (VVF).

### Human embryonic kidney 293T cells and transfection

HEK293T cells were routinely cultured in Dulbecco’s modified Eagle Medium (DMEM, Thermofisher scientific) and 10% foetal calf serum. Cells were periodically split using Versene solution (Gibco) and mechanical dissociation. Dissociated cells were seeded into 6 well plates and when cells reached 70% confluence, were transfected using JetPEI following the manufacturer’s protocol (PolyPlus transfection). A total of 3 μg of DNA/per 35 cm well was combined with NaCl and JetPEI and after 20 mins added to HEK293T cells. The next day transfected cells were re-plated onto cover slips and used 24 hrs later.

### DRG neuron culture and electroporation

Briefly, mice were sacrificed and spinal columns removed. Dorsal root ganglia were rapidly dissected and enzymatically digested at 37°C for 60-90 mins in dispase type II (4.7 mg/ml) and collagenase type II (4 mg/ml). Cells were briefly centrifuged and HBSS/CollagenaseDispase removed. Pre-warmed culture media (Neurobasal, 2% B-27 supplement, 1% Penicillin streptomyocin) was added and cells were mechanically dissociated using fire-polished pipettes. Neurons were transfected via electroporation using the Neon system (Life technologies). Dissociated cells were re-suspended in 10 μl of Buffer R plus 1 μg of total plasmid DNA per 50-100,000 cells. The electrical protocol applied was three 1500-V pulses of 10 ms duration. Cells were immediately plated on Poly-D-lysine/Laminin coated cover slips with the addition of growth factors (mouse nerve growth factor (50 ng/ml; NGF, PeproTech) and 10 ng/ml glial-derived neurotrophic factor (GDNF, PeproTech)). Cells were used for further experiments up until day 4 *in vitro*.

### Generation and culture of induced pluripotent stem cells

Healthy control iPSCs, AD2-1 and AD3-1 (StemBANCC Consortium), were derived from fibroblasts as described previously (Clark et al., 2017). Data obtained AD2-1 and AD3-1 hiPSCs were pooled and used as control. Another line, RCi002-A, was derived from a patient with inherited erythromelalgia and carries the F1449V mutation in SCN9A (EBiSC Consortium) (*43*). All lines were separately reprogrammed by non-integrating Sendai viral vectors using the CytoTune-iPS Reprogramming Kit (ThermoFisher). For quality control all iPSC lines were subject to strict checks before initiation of differentiation, including; tests for Sendai virus clearance, FACS for pluripotency markers, genomic integrity checks, cytoSNP analysis for copy number variation and embryoid body tri-lineage differentiation experiments. Cells are also confirmed as negative for Mycoplasma before cryopreservation. iPSCs were maintained in mTesR1 (StemCell Technologies) or StemFlex (Life Technologies) on Matrigel (Corning) coated dishes. Cells were routinely passaged at 80% confluence with EDTA (Life Technologies). Medium was supplemented with Y-27632 (Tocris) when thawing iPSCs.

### Differentiation of human induced pluripotent stem cells to sensory neurons

Human iPSCs were differentiated following the Chamber’s protocol, with modifications (Clark et al 2017, 2021). In brief, cells were passaged using Versene EDTA (ThermoFisher) and plated at high density. Neural induction commenced with the addition of SMAD inhibitors SB431542 (Sigma, 10 mM) and LDN-193189 (Sigma, 100 nM) to KSR base medium (Knockout-DMEM, 15% knockout-serum replacement, 1% Glutamax, 1% nonessential amino acids, 100 mM b-mercaptoethanol, (ThermoFisher)). Three additional small molecules were introduced on day 3 (CHIR99021 (Sigma, 3 mM), SU5402 (Sigma, 10 mM) and DAPT (Sigma, 10 mM). The dual SMAD inhibitors were withdrawn on day 5. The base medium was gradually transitioned to N2/B27 medium (Neurobasal medium, 2% B27 supplement, 1% N2 supplement, 1% Glutamax, (ThermoFisher)) in 25% increments. Cells were replated onto glass coverslips at day 12 of the differentiation in N2/B27 medium supplemented with four recombinant growth factors at 25ng/ml (BDNF; ThermoFisher, NT3, NGF, GDNF; Peprotech). CHIR90221 was included for 4 further days. Medium changes were performed twice weekly after replating onto coverslips. If required, Cytosine b-D-arabinofuranoside (araC, 1-2 mM, Sigma) was included in the medium soon after replating to kill the few non-neuronal dividing cells remaining in the culture. AraC was withdrawn from the medium once a pure neuronal culture was obtained, as judged by the absence of morphologically non-neuronal cells on phase-contrast light microscopy. This state was typically achieved 2-3 weeks after replating. From day 28, the concentration of all four recombinant growth factors was reduced to 10ng/ml. Phenol-free Matrigel (Corning, 1:500 dilution) was included in all medium changes from day 28 onward. Medium changes were performed twice weekly. AAVs (AAV9-CAG-mCherry-T-PSAM^4^-GlyR, multiplicity of infection (MOI): 1M, and AAV9-CAG-mCherry, MOI: 100K) were added to the cultures around day 50. AAVs remained in culture for 7 days without media change. Biweekly media changes resumed thereafter. Cells were used for experiments at least 4-6 weeks post AAV infection.

### Whole-cell patch clamp recordings

Voltage-clamp recordings using an Axopatch 200B amplifier and Digidata 1550 acquisition system (Molecular Devices) were performed at room temperature. Data were sampled at 20kHz and low-pass filtered at 5 kHz. Series resistance was compensated 70% –85% to reduce voltage errors. All data were analyzed by Clampfit 10 software (Molecular Devices). mCherry+ HEK cells, DRG neurons and hiPSC-sensory neurons, were detected with an Olympus microscope with an inbuilt Cy3 or Texas red filter set. Borosilicate glass capillaries (1.5 mm OD, 0.84 mm ID; World Precision Instruments) were pulled to form patch pipettes of 2–5 MΩ tip resistance and filled with an internal solution containing (mM): 100 K-gluconate, 28 KCl, 1 MgCl2, 5 MgATP, 10 HEPES, and 0.5 EGTA; pH was adjusted to 7.3 with KOH and osmolarity set at 305-310 mOsm (using glucose). Cells were maintained in a chamber constantly perfused with a physiological extracellular buffer containing (mM): 140 NaCl, 4.7 KCl, 2.5 CaCl2, 1.2 MgCl2, 10 HEPES and 10 glucose; pH was adjusted to 7.4 with NaOH and osmolarity set at 310-315 mOsm (using glucose). uPSEM^792^ and varenicline fresh daily, diluted in extracellular buffer to a final concentration of 10 nM and 20 nM respectively, unless otherwise stated. Agonists were delivered to the cells via a perfusion system. All post agonist recordings were made 15 mins post application. Membrane conductance was measured in voltage-clamp mode with a 100ms voltage ramp from -90 mV to +40 mV every 10 s. Liquid junction potential (−13 mV) was corrected. The resultant linear current gradient was used to calculate membrane conductance using the rearranged Ohm’s law equation where V = voltage, I = current, R =resistance, C = conductance. V=IR, C=1/R, C=I/V. Resting membrane potential (RMP) was measured in bridge mode (I=0). In current-clamp mode, neurons were held at -60 mV. Input resistance (Rin) was derived by measuring the membrane deflection caused by a 20 pA current step. Rheobase was determined by applying 50 ms depolarising currents of increasing steps of 25 pA until action potential (AP) generation.

### Cell-attached patch clamp recordings

Cell-attached patch clamp was performed at 32°C using the same experimental set up as above and using the same solutions. Once a Giga-seal was achieved, cell-attached recordings were sampled in voltage clamp. The spontaneous activity of each cell was recorded for 5-10 mins. A spontaneously active cell was defined as firing at least 1 AP in 5 mins. Neurons were recorded in the presence of 20 nM varenicline throughout the duration of the experiment.

### Spinal cord slice recordings

Adult mice, that had received a subcutaneous viral injection when neonates, were anaesthetized with ketamine/xylazine (i.p. 90mg/kg / 10mg/kg), and perfused transcardially with ice-cold oxygenated (95% O2, 5% CO2) sucrose-based artificial cerebrospinal fluid (sACSF) containing (in mM): 100 sucrose, 63 NaCl, 2.5 KCl, 1.2 MgCl2, 1.2 NaH2PO4, 25 NaHCO3, 25 glucose and 1 kynurenate. Spinal cords were carefully obtained by laminectomy in ice-cold sACSF with dorsal roots attached. Parasagittal slices (300 μm) were cut on a vibratome (Leica VT 1200) in ice-cold sACSF. Slices were then transferred to a submerged chamber containing oxygenated NMDG-based recovery ACSF (rACSF) for 15 minutes at 34 °C, containing (in mM): 93 NMDG, 2.5 KCl, 1.2 NaH2PO4, 30 NaHCO3, 20 HEPES, 25 Glucose, 5 Na ascorbate, 2 thiourea, 3 Na pyruvate, 10 MgSO4 and 0.5 CaCl2, and adjusted to pH 7.4 with HCl. After recovery incubation, slices were transferred to oxygenated ACSF where they were maintained at room temperature prior to transfer to the recording chamber. ACSF was composed of (in mM): 126 NaCl, 2.5 KCl, 2 MgCl2, 2 CaCl2, 1.25 NaH2PO4, 26 NaHCO3 and 10 glucose. Neurons were visually identified in an Olympus BX51 microscope equipped with infrared differential interference contrast (IR-DIC) and 40x water-immersion-objectives. Patch pipettes (5–7 MΩ) were pulled on a horizontal puller (P-1000; Sutter) and filled with the following intracellular solution (in mM): 135 K-gluconate, 5 KCl, 2 MgCl2, 10 HEPES, 4 ATP-Na, 0.4 GTP-Na, 0.1% Lucifer-Yellow (LY, Sigma), pH 7.3 adjusted with KOH. A suction electrode filled with ACSF was placed in a dorsal root was to deliver electrical stimulation. Signals were amplified with a Axopatch 200B amplifier (Molecular Devices), digitized with a Digidata 1440 (Molecular Devices), and recorded using pClamp 10 software (Molecular Devices). Data were filtered at 5 kHz and sampled at 10 kHz. Neurons were maintained at -70 mV (corrected for liquid junction potential of -10 mV). Monosynaptic responses were determined by the absence of failures in a 10 Hz train of 10 electrical stimuli. Agonists were superfused onto slices at a rate of 2 ml/min in normal oxygenated ACSF. Only neurons with a resting potential more negative than -50 mV and stable access resistance (<25 MΩ) during the recording were included for subsequent analysis.

### Pup subcutaneous injection

Neonatal (P5-P7) mice were briefly removed from their homecage and injected with 10 μl of virus, subcutaneously at the nape of the neck. They were rubbed in their bedding before being returned to their home cage. Mice were used for tissue or electrophysiology at least 4 weeks later.

### Intrathecal infusion

This procedure has been described in detail previously (*38*). Briefly, each animal was anaesthetised using 2% isoflurane and prepared for surgery by shaving a region over the thoracic vertebrae. T-10 and T-11 vertebrae were located, an incision was made followed by removal of soft tissue to expose the dura and spinal cord. A drop of lidocaine was applied to the dura for approximately 1-2 mins then removed. Using a 30 gauge needle the dura was carefully punctured (CSF leak at this point suggested a successful puncture). A cannula system was designed by connecting tubing of decreasing size until the final cannula tip measured 0.008 in (O.D) x 0.004 in (I.D). The end of the cannula was inserted approximately 1 cm caudal into the subdural space. Using a syringe pump driver, 8 μl of AAV was injected into the subdural space at a rate of 1 μl/min. Following injection, the cannula was allowed to rest in position for 2 min before being slowly removed. The dura was coated with a single drop of dura gel (Cambridge NeuroCare) to seal the dura and prevent further CSF leak. Finally, the incision site was sutured closed and appropriate post-operative care and analgesics given (local 2 mg/kg Marcain, AstraZeneca and systemic 5 mg/kg Rimadyl, Pfizer). Animals were used for behaviour or histology at least 6 weeks post-surgery.

### Intra-articular injection

Intra-articular injections of AAVPHP.s-eGFP or AAVPHP.s-mCherry-T-PSAM^4^-GlyR were made to both knees under anaesthesia (100 mg/kg ketamine and 10 mg/kg xylazine, delivered intraperitoneally) when mice were aged 6 weeks. 4-weeks later, after capturing baseline behaviours, mice were anaesthetised and one knee (side determined randomly) received an intra-articular injection of 10 µg complete Freund’s adjuvant (CFA; Chondrex) to induce inflammation. The width of each knee joint was measured with digital callipers before and 24-hours post-CFA injection. After assessing post-CFA behaviour, mice received an intraperitoneal injection of 0.3 mg/kg varenicline (from a 0.06 mg/ml stock).

### Chemogenetic agonists

uPSEM^792^(*31*) (Tocris Bioscience) the designer PSAM^4^-GlyR agonist, was reconstituted in saline and injected i.p. at a final concentration of 5mg/kg (unless otherwise stated). Varenicline (Merck, PZ0004) the clinically relevant PSAM^4^-GlyR agonist, was reconstituted in saline and injected i.p. at a final concentration of 0.3mg/kg.

### Behavioural analysis related to the hind paw

Both male and female mice were used in this study, and mice were tested at a consistent time of day, in the same environment by the same experimenter. Mice were habituated to their testing environment and equipment prior to behavioural test days. Mice we tested in a random order and randomly assigned a test box each day. The experimenter was blind to the animal group until after the behavioural analysis was complete. Mice were tested on three different days to obtain an average baseline value. Except in the case of injury, all sensory threshold data represent the average of right and left hind paws. Mice were tested after (1-2hrs unless otherwise stated) receiving i.p. uPSEM^792^ or varenicline. Mice were re-tested at least 24 hrs later for washout experiments.

### von Frey

Mice were elevated on a wire mesh base in a test box (5 × 5 × 10 cm), and acclimatized to the equipment for 30–60 min. The plantar hind paws were tested using calibrated von Frey hairs (Linton Instrumentation) using the ‘up-down’ method (Dixson 1980) to evaluate their 50% paw withdrawal thresholds. For spared nerve injury experiments mice were test for mechanical sensitivity over the course of the injury (days 6, 7, 14, 21, 28).

### Brush

The plantar hind paws of mice were brushed (1 cm s-1) with a fine artists paint brush. Each mouse received 5 successive stimuli on alternate hind paws (10s apart), twice. The number of responses were recorded. A response included, lifting, flicking or moving the hind paw or walking away from the stimulus.

### Pinprick

Noxious mechanosensation was assessed by the pinprick test described previously (*79*) Mice were housed and acclimatized similarly to the von Frey test. Mice were tested on their plantar hind paws using a sharp pin attached to a 1 g calibrated von Frey filament. Mice were video recorded using a GoPro at 240 fps, and the latency to withdraw from the pinprick analysed by an investigator blind to treatment groups. Three measurements were taken for each hind paw per trial and plotted latency represents the average of both paws.

### Hargreaves

Thermal thresholds were assessed using an infrared light source applied to the plantar surface of each hind paw. Three measurements were taken for each hind paw and the averaged latency to withdraw was measured.

### Dry Ice

Noxious cold thresholds were measured using the dry ice assay. Mice were elevated on a glass platform in a test box. Pieces of dry ice were place into a 2 ml syringe (top cut off). The syringe filled with dry ice was placed against the glass from below (where hind paws were flat and visible). Latency to withdraw paws form the dry ice/glass was measured. Three measurements were taken for each hind paw

### Beam task

The Beam test (*80*) apparatus consisted of a 1-m long horizontal beam of 12-mm width suspended from the ground. Mice were placed at one end of the beam next to a light source and walked across the beam towards a darkened box/house. Trials were recorded once per day and video footage was used by an investigator blind to treatment groups, to assess number of steps and number of missteps. Baseline data were generated from the average of two trials on separate days. 1 mouse was excluded from the beam test due to abdominal obesity, which prevented the hind paws from being able to reach the beam.

### Behavioural analysis related to the knee

All behavioural experiments were conducted in the presence of one female and one male investigator. Mice were allowed 30 minutes to acclimatise to the testing room before commencing behavioural assays. Both male and female mice were used in this study. Baseline behaviours were tested 2 hours prior to i.a. CFA injection. Mice were tested again 24 hrs post-CFA and 1-2 hrs post-varenicline, mice were tested in a random order each time. The experimenters were blind to the animal group until after the behavioural analysis was complete.

### Digging

The digging behaviour of mice was measured as a readout of spontaneous pain. Testing involved placing individual mice into standard 49 × 10 × 12 cm cages filled with ∼4 cm tightly packed fine-grain aspen midi wood chip bedding substrate (LBS Biotechnology). Mice were allowed 3 minutes to explore testing cages under video surveillance. Training sessions were carried out the day before baseline behaviours were captured, during these sessions mice were placed in test cages as per a normal test, however, mice that did not dig for at least 15 seconds were subsequently placed in a test cage with a cage-mate that did meet this criterion until both animals demonstrated digging behaviour. Following test digs, the number of visible burrows at the end of the 3 minutes was recorded. Digging duration was scored independently by two investigators following the conclusion of each study and blinding of the acquired videos; since the scores of investigators was well correlated (R2 = 0.86, across 102 videos) an average is reported as the digging duration.

### Rotarod

Locomotor function and coordination was assessed using a rotarod (Ugo Basile). Mice were placed on the rotarod at a constant speed (7 rpm) for 1 minute before starting an accelerating programme (7-40 rpm, over 5 minutes), test runs were video recorded. Mice were removed from the rotarod if they fell, following two consecutive passive rotations or after 6 minutes of the accelerating program, whichever occurred first. Mice were first placed on the rotarod the day before baseline behaviours were captured to gain some familiarity with the assay. The latency to passive rotation or fall was timed by 1 investigator at the conclusion of each study following blinding of the acquired videos.

### Pressure Application Measurement

Mechanical sensitivity of the knee joint was assessed using a pressure application measurement device (Ugo Basile). Mice received no training in this assay before acquiring baseline sensitivity and digging, and rotarod behaviours were always assessed before the application of pressure to the knee joint. Animals were scruffed before the force transducer was used to apply gradual force to each of the animals’ knee joints, by squeezing the joint medially. The withdrawal threshold was recorded when an animal withdrew the limb being tested, or after 450 g force was applied, whichever occurred first. Each animal was tested twice per time point, with a short break between tests, withdrawal force is reported as an average of the two measurements taken at each time point.

### Chemical pain model - formalin assay

Mice were chosen at random from their home cage and the left hind paw of each mouse was injected subcutaneously with 2% formalin and mice were immediately placed in a test box (5 x 5 x 10 cm), on a glass base, which was elevated above a camera. The perimeter and roof of the test box consisted of mirrors to allow good visualisation of the injected hindpaw. The mice were video recorded in this environment for 1 hr while the experimenter left the room. Off line analysis was used to measure nocifensive behaviours of the injected hind paw (lifting, licking, flinching, shaking) every 5 mins for 60 mins. The formalin assay was also analysed in 2 phases; 1st phase 0-15 mins, 2nd phase 15-60 mins. All formalin behaviour was conducted 1 hrs post 0.3mg/kg i.p. of varenicline.

### Neuropathic pain model

For the spared nerve injury model we ligated and transected the sural and common peroneal branches of the sciatic nerve, leaving the tibial nerve intact (tSNI). Animals received postoperative analgesia as detailed above and were assessed behaviourally from 6 days post-SNI. Any animals that did not show a reduction in mechanical thresholds by 7 days post-SNI were excluded from further assessment and were not included in the final analysis. The lack of mechanical hypersensitivity led us to exclude 1/24 animals.

### Immunohistochemistry

Animals were deeply anaesthetized with pentobarbital and the blood cleared from all tissues by perfusing saline through the vascular system. Mice were then perfuse-fixed using 4% paraformaldehyde (PFA). Tissues were then collected and post-fixed in 4% PFA accordingly (DRG: 1–2 h, spinal cord: 24 h). All tissues were cryoprotected in 30% sucrose for a minimum of 48 h, followed by embedding the tissue and sectioning on a cryostat. (DRG: 12 μm, spinal cord: 15 μm). Cultured cells were fixed with 4% PFA for 10 min and treated similarly to other tissues. Standard immunohistochemistry protocols were used. Briefly, fixed/sectioned samples were washed in PBS and blocked in a blocking solution (5% normal donkey serum, 0.3% TritonX-100, PBS) for 1hr at room temperature (RT). Primary antibodies (see Table below) were diluted in blocking solution and applied to tissue or cells overnight at RT. The next day samples were washed in a wash solution (0.3% TritonX100, PBS) followed by a 2hr incubation with secondary antibodies diluted in wash solution at RT. Samples were mounted using Vectorshield and imaged on a confocal microscope (Zeiss LSM-710). Images were analysed using Fuji/ImageJ (NIH). For quantification at least three sections per animal were used, with at least 3 animals per group.

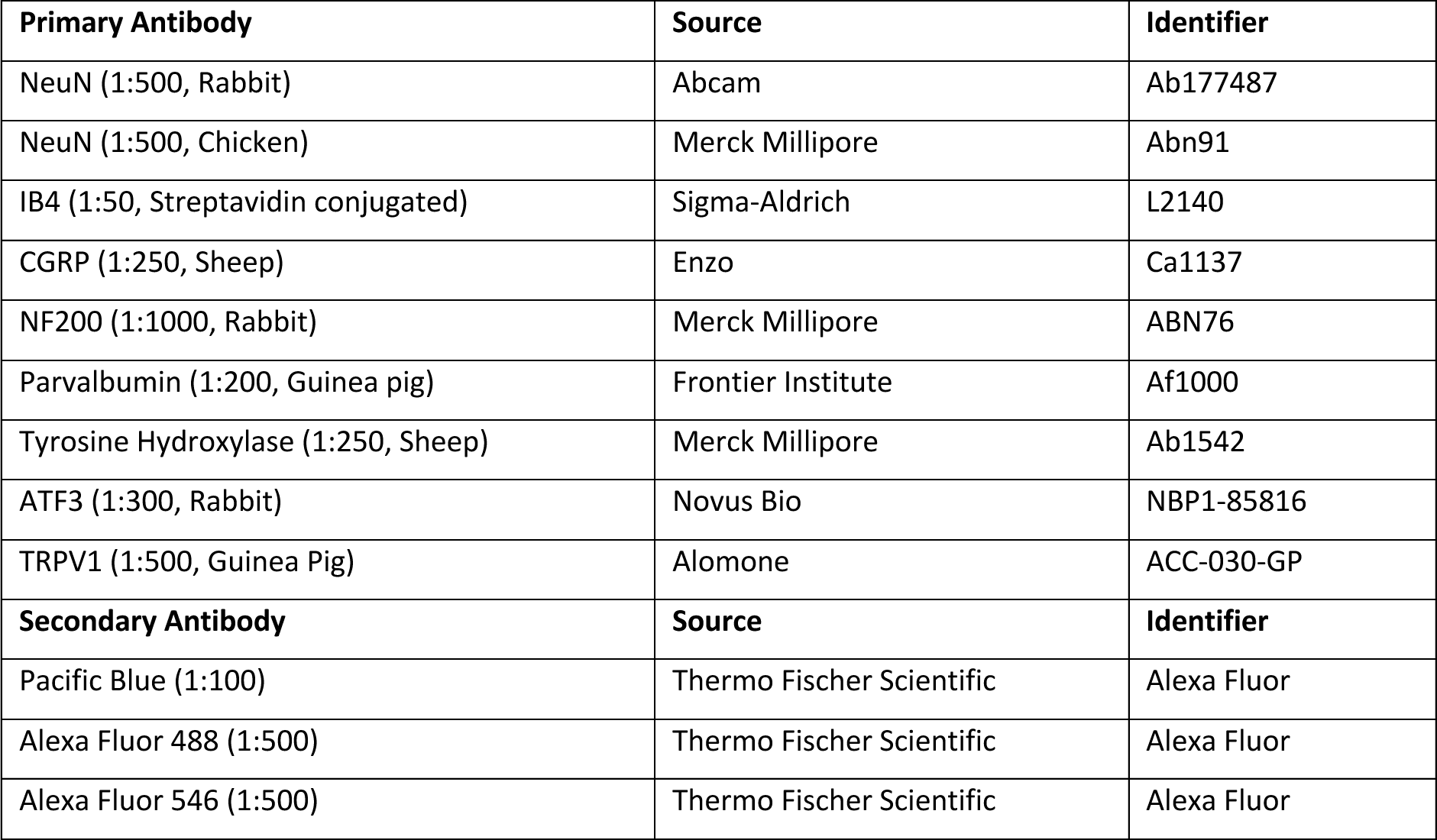

| **Primary Antibody** | **Source** | **Identifier** |
| --- | --- | --- |
| NeuN (1:500, Rabbit) | Abcam | Ab177487 |
| NeuN (1:500, Chicken) | Merck Millipore | Abn91 |
| IB4 (1:50, Streptavidin conjugated) | Sigma-Aldrich | L2140 |
| CGRP (1:250, Sheep) | Enzo | Ca1137 |
| NF200 (1:1000, Rabbit) | Merck Millipore | ABN76 |
| Parvalbumin (1:200, Guinea pig) | Frontier Institute | Af1000 |
| Tyrosine Hydroxylase (1:250, Sheep) | Merck Millipore | Ab1542 |
| ATF3 (1:300, Rabbit) | Novus Bio | NBP1-85816 |
| TRPV1 (1:500, Guinea Pig) | Alomone | ACC-030-GP |
| **Secondary Antibody** | **Source** | **Identifier** |
| Pacific Blue (1:100) | Thermo Fischer Scientific | Alexa Fluor |
| Alexa Fluor 488 (1:500) | Thermo Fischer Scientific | Alexa Fluor |
| Alexa Fluor 546 (1:500) | Thermo Fischer Scientific | Alexa Fluor |

### Sample sizes and statistical analysis

Behavioural sample sizes were calculated using the software G*power2 with p-values of 0.05 and a power of >0.8. In pain behaviour outcomes the effect size is taken as a 30% difference which parallels what is often seen as clinically relevant changes in pain ratings in human. Data on variance was generated from our provisional and published data using these outcome measures. The primary outcome was evoked behaviour, experimental unit was mouse, units per group was 9 biological replicates, 3 measurements, effect size 30 (f=0.6), and RM ANOVA was chosen for the primary outcome statistical test. Note that for group sizes in cohorts that underwent AAV intrathecal injection n of 1 was added to this sample size in order to deal with surgical attrition. If mice underwent spared nerve injury n of 1 was also added to in order to deal with any surgical attrition. All data was tested for normality using the D’Agostino-Pearson normality test and the appropriate parametric or non-parametric statistical tests used accordingly. All statistical tests used were two-tailed. Statistical comparisons were made using a Student’s t-test or Mann Whitney U-test. In experimental groups in which multiple comparisons and repeated measures were made two-way analysis of variance (ANOVA) tests with appropriate post-hoc tests were performed. For electrophysiological data, n equals number of cells from at least 3 animals All data is represented as mean ± the standard error of the mean (S.E.M.) unless otherwise stated. Statistical significance is indicated as follows * P < 0.05, ** P < 0.01, *** P < 0.001, **** P <0.0001. The statistical test used is reported in the appropriate figure legend. Graph Pad prism 9 was used to perform statistical tests and graph data. Adobe illustrator CS5 was used to create schematics and medical graphics were obtained from Smart servier free medical art (smart.servier.com).

## Acknowledgements

**Acknowledgments:** For the purpose of open access, the author(s) has applied a CC BY public copyright licence to any Author Accepted Manuscript version arising from this submission.

**Funding:** This research was funded in whole, or in part, by the Wellcome Trust (102645/Z/13/Z, 202747/Z/16/Z, and 109116/Z/15/Z). D.L.B. is a Wellcome Senior Investigator (223149/Z/21/Z). D.L.B. and S.J.M. acknowledge funding from the UK Medical Research Council (grant ref. MR/T020113/1). H.Hu. acknowledges the funding from China Scholarship Council (CSC). E.St.J.S. and L.A.P. acknowledge funding from the MRC (MR/W002426/1). H.Hi. was funded by a BBSRC/GSK iCASE PhD studentship (BB/V509528/1)

## Author contributions

Conceptualization: JPS, SJM, EStJS, DLB

Methodology: JPS, SJM, LAP, HHi, MAA, SRZ, MBR, HHu, XY, AJC, EStJS, DLB

Investigation: JPS, SJM, LAP, HHi, MAA, SRZ, MBR, HHu, XY, AJC

Visualization: JPS, SJM, LAP, HHi

Funding acquisition: EStJS, DLB.

Project administration: JPS., SJM

Supervision: SJM, EStJS, DLB

Writing – original draft: JPS., SJM, DLB

Writing – review & editing: JPS, SJM, LAP, HHi, MAA, SRZ, MBR, HHu, XY, AJC, EStJS, DLB

**Competing interests:** D.L.B. has acted as a consultant on behalf of Oxford Innovation for Abide, Amgen, G Mitsubishi, Tanabe, GSK, TEVA, Biogen, Lilly, Orion and Theranexus.

**Data and materials availability:** All data and materials will be made available upon reasonable request to the corresponding author(s). Construct(s) generated in this study will be made available on Addgene.

**Fig. S1.**
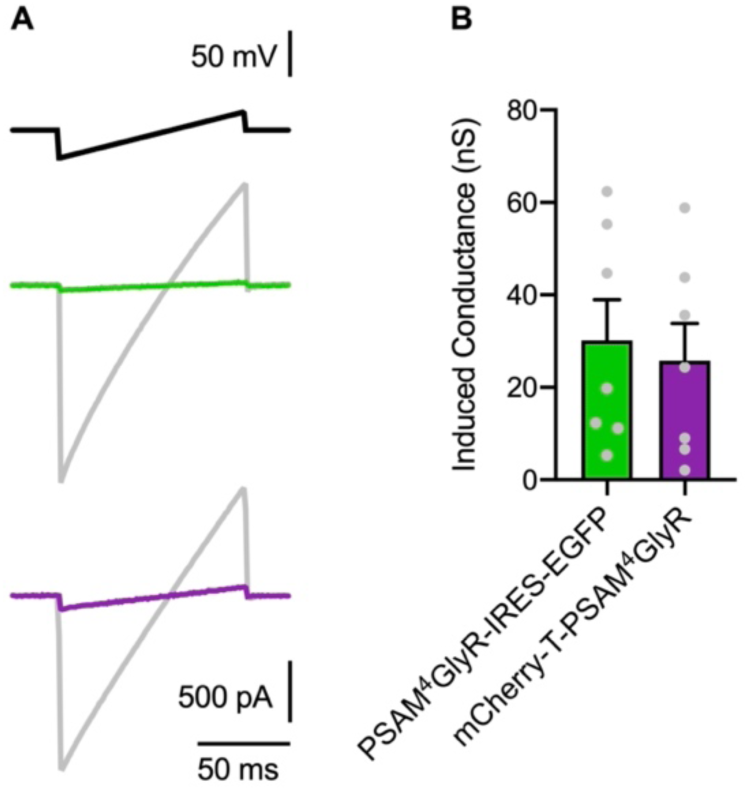
Modification of the construct containing PSAM^4^-GlyR does not affect function. **(A)** Representative traces showing increased membrane conductance after varenicline administration in HEK293t cells transduced with PSAM^4^-GlyR-IRES-GFP (green) and mCherry-T-PSAM^4^-GlyR (purple). **(B)** Quantification of induced membrane conductance in HEK293t cells after varenicline administration (Unpaired t-test, P > 0.05). Data expressed as mean ± S.E.M.

**Fig. S2.**
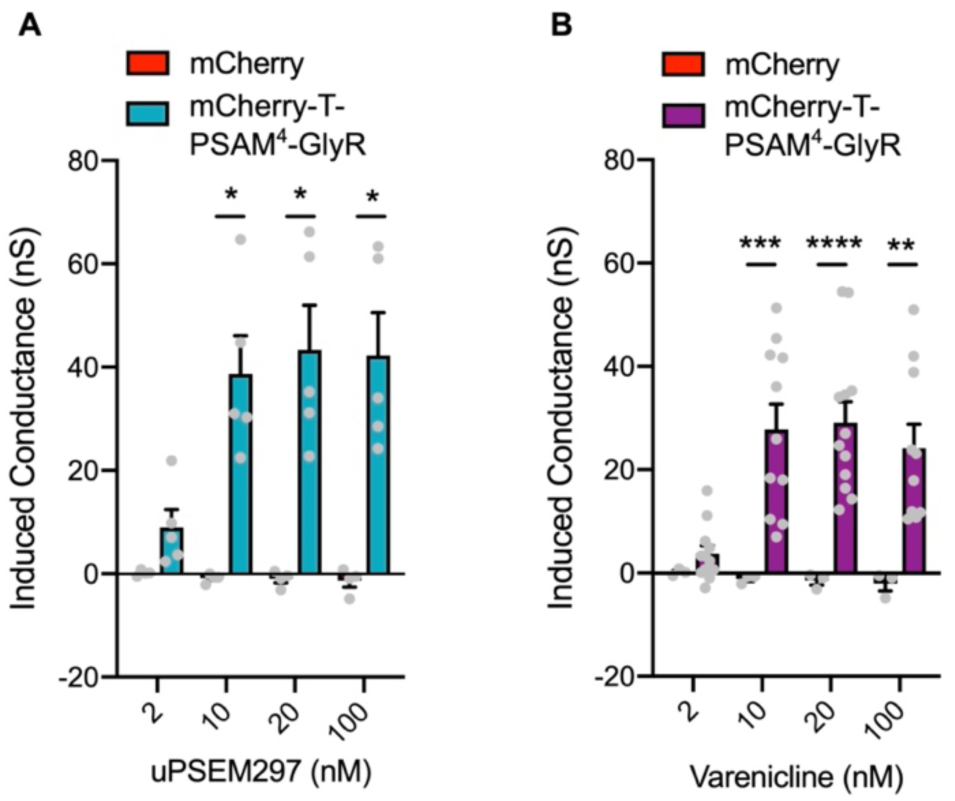
Dosage of PSAM^4^-GlyR agonist required for increase in membrane conductance. **(A)** Quantification of changes in membrane conductance by application of 2, 10, 20 and 100 nM uPSEM^792^ in dissociated sensory neurons transduced with mCherry and mCherry-T-PSAM^4^-GlyR (RM-two way ANOVA, post-hoc Bonferroni test, ** P = 0.001, *** P = 0.0005, **** P < 0.0001). **(B)** Summary of increase in membrane conductance after application of 2, 10, 20 and 100 nM varenicline in dissociated sensory neurons transduced with mCherry and mCherry-T-PSAM^4^-GlyR (RM-two way ANOVA, post-hoc Bonferroni test, * P < 0.05). Data expressed as mean ± S.E.M.

**Fig. S3.**
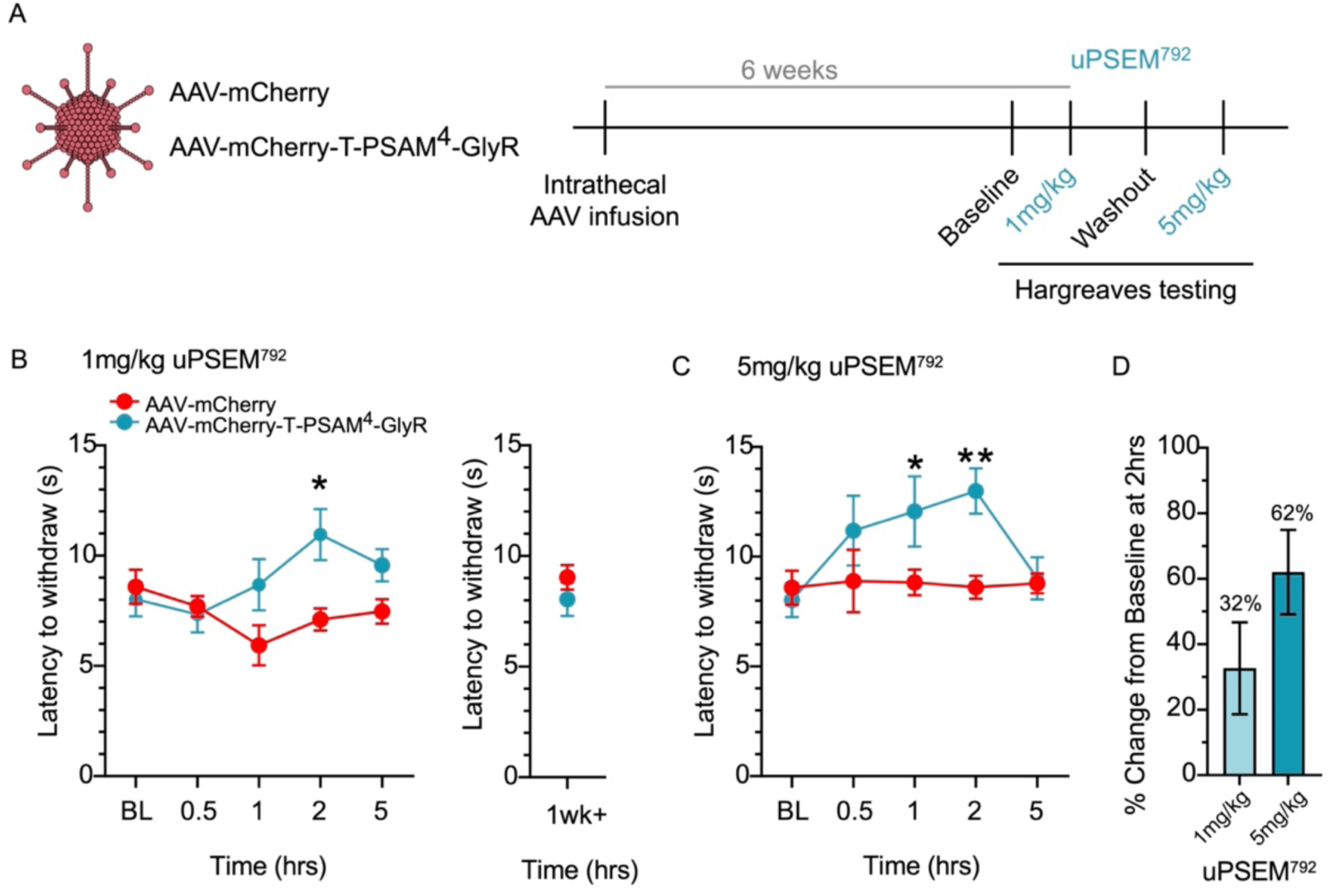
uPSEM^792^ dosage and time-course of agonist induced PSAM^4^-GlyR silencing of thermal nociception. (**A**) Schematic of the experimental timeline, AAVs were delivered intrathecally and 6 weeks later Hargreaves testing was conducted post i.p. of 1mg/kg or 5mg/kg uPSEM^792^. (**B**) The latency to withdraw from a noxious radiant heat source was measured 0.5, 1, 2, and 5 hrs post 1mg/kg uPSEM^792^. AAV-mCherry-T-PSAM^4^-GlyR mice were significantly hyposensitive to the Hargreaves apparatus 2 hrs post 1mg/kg uPSEM^792^ compared to AAV-mCherry mice. (mCherry: n = 6 mice, PSAM^4^-GlyR: n = 8 mice, RM-two way ANOVA, post-hoc Bonferroni test, * P = 0.013). Behavioural hyposensitivity returned to normal after 5 hrs and remained normal for 1 week. (**C**) AAV-mCherry-T-PSAM^4^-GlyR mice were significantly hyposensitive to the radiant heat source, 1 and 2 hrs post 5mg/kg uPSEM^792^ compared to AAV-mCherry mice (mCherry: n = 3 mice, PSAM^4^-GlyR: n = 4 mice, RM-two way ANOVA, post-hoc Bonferroni test, * P < 0.05, ** P < 0.01). (**D**) The percentage change in withdrawal latency at 2hrs (from baseline) for 1mg/kg (n = 8 mice) and 5mg/kg uPSEM^792^ (n = 4 mice). Data expressed as mean ± S.E.M.

**Fig. S4.**
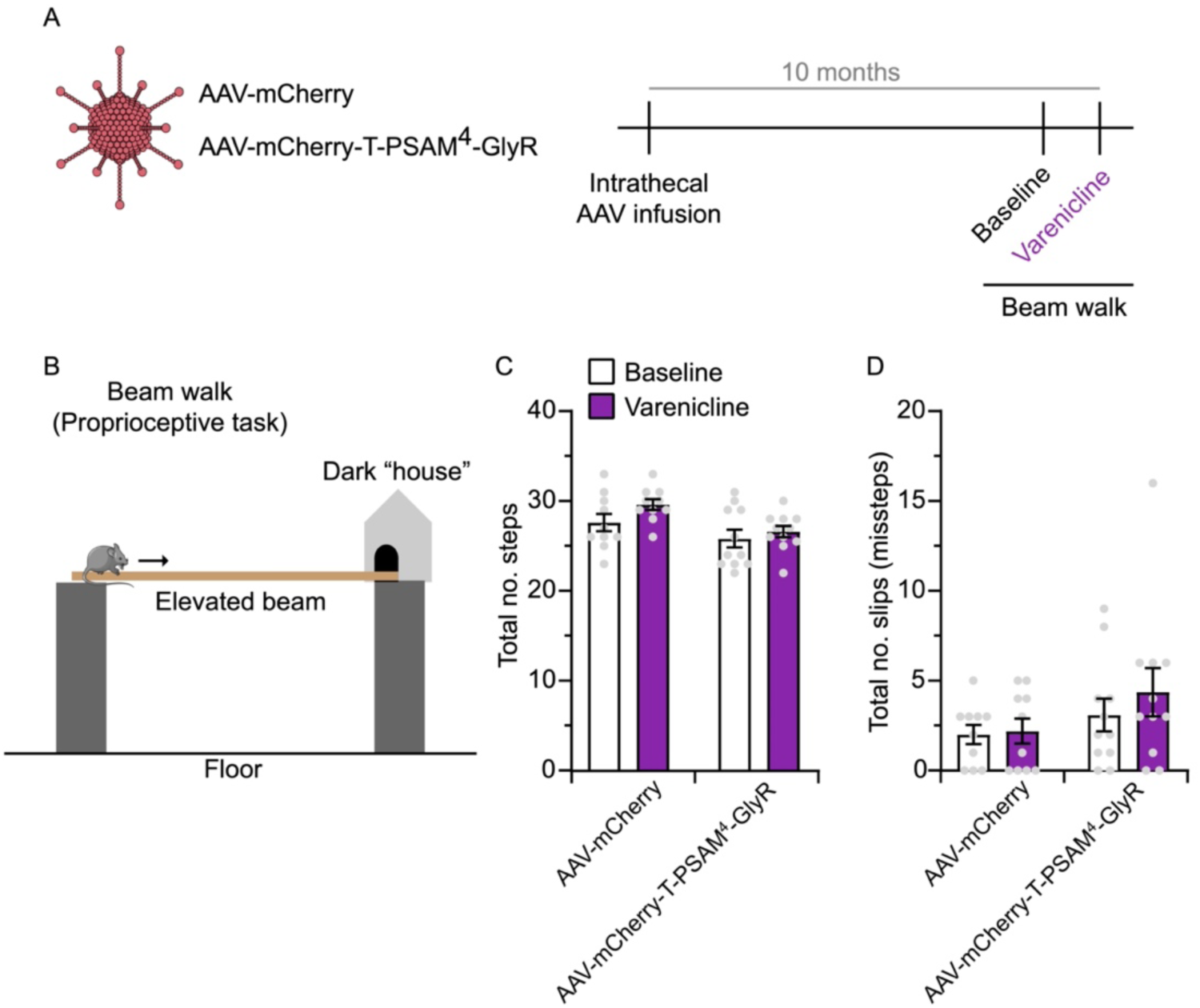
Proprioceptive behaviours are preserved following PSAM^4^-GlyR mediated silencing of sensory neurons. (**A**) Schematic of the experimental design. (**B**) Depiction of the beam task, mice are trained to walk along a thin beam toward a dark house/space. Steps along the beam are video recorded. (**C**) Quantification of total number of steps during the task, which was unchanged between groups or following varenicline administration. (**D**) Quantification of the total number of slips or mistakes during the task. The number of mistakes was unchanged in both groups following varenicline (mCherry: n = 10 mice, PSAM^4^-GlyR: n = 11 mice, all data sets RM-two way ANOVA, post-hoc Bonferroni test, P > 0.05). Data expressed as mean ± S.E.M.

**Fig. S5.**
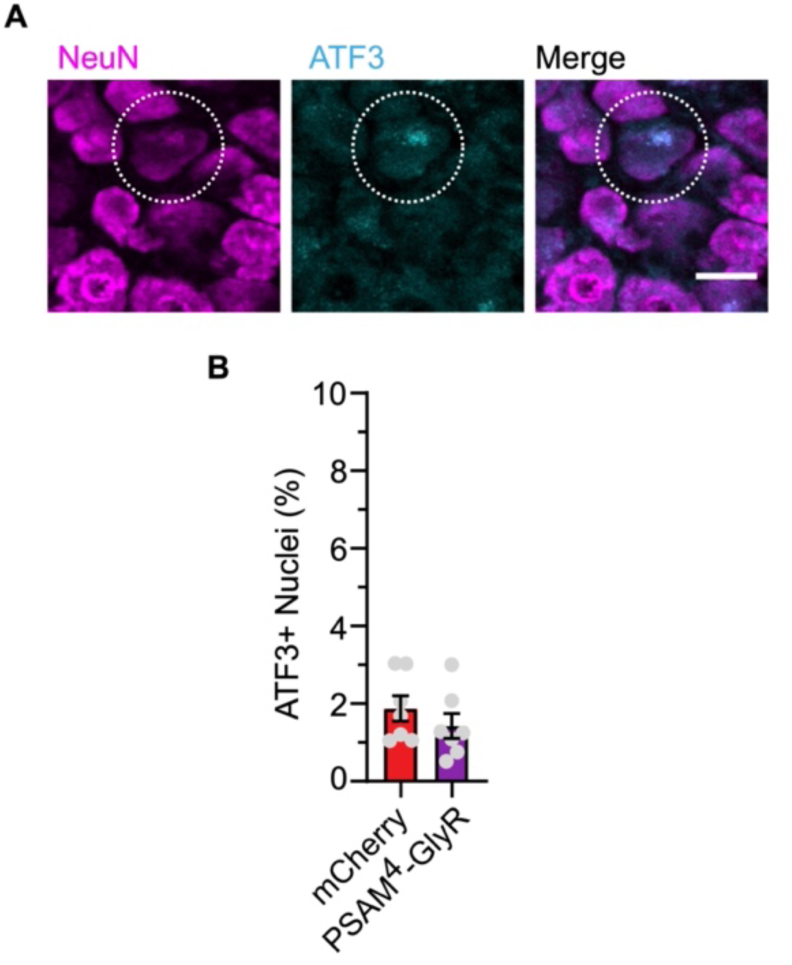
Long-term expression of mCherry-T-PSAM^4^-GlyR, does not result in up regulation of the injury marker ATF3. (**A**) Example image of a DRG neuron with an ATF3 positive nucleus (scale bar 25 um). (**B**) Quantification of the percentage of ATF3 + nuclei in DRG neurons from mice that received AAV-mCherry or AAV-mCherry-T-PSAM^4^-GlyR (mCherry: n = 7 mice, 19/1014 neurons, PSAM^4^-GlyR: n = 7 mice, 16/1324 neurons). Data expressed as mean ± S.E.M. n.s. P > 0.05, calculated by an unpaired t-test.

**Fig. S6.**
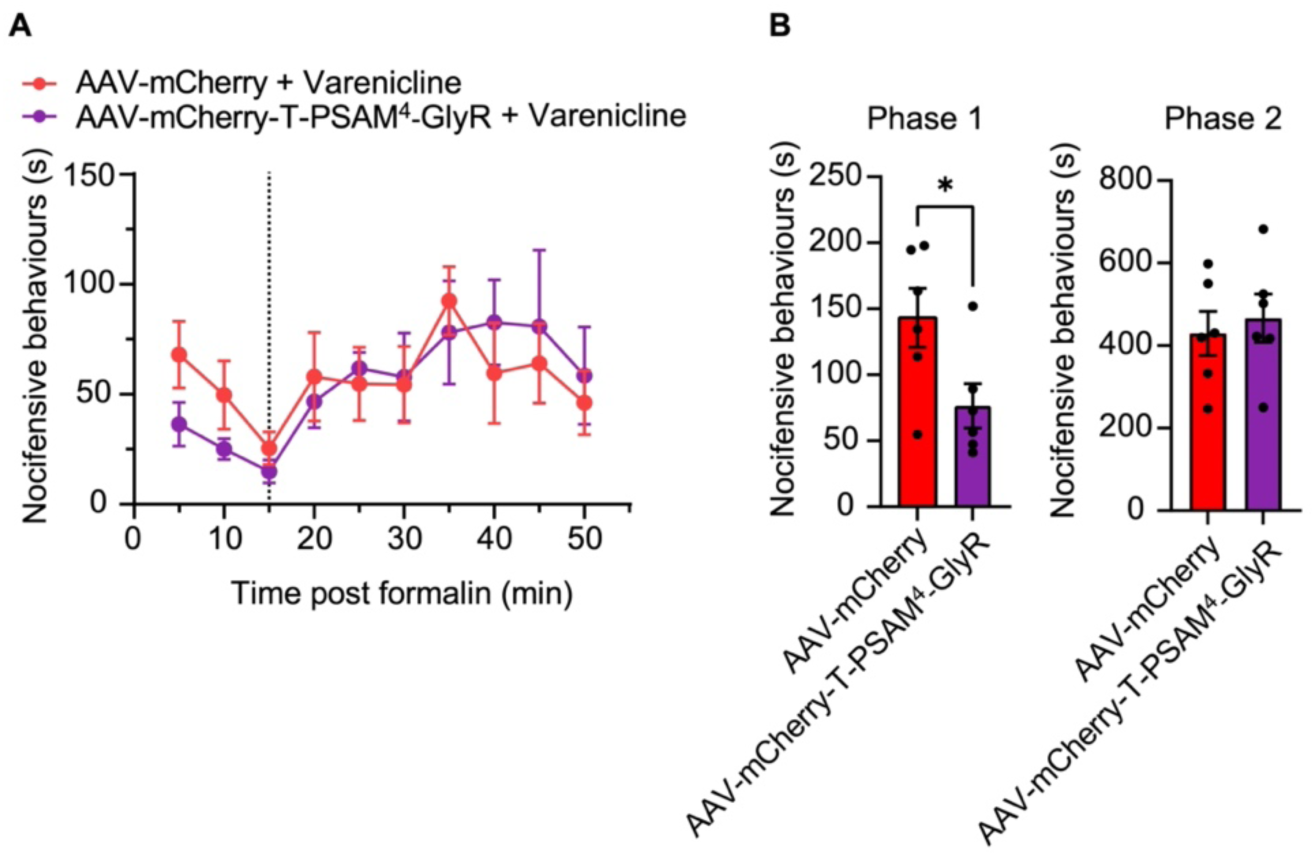
PSAM^4^-GlyR chemogenetic silencing of primary afferents reduces chemical-induced pain. (**A**) AAV-mCherry or AAV-mCherry-T-PSAM^4^-GlyR mice were both given varenicline and 1 hr later received an injection of 2% formalin in the hind paw. Nocifensive behaviours over the following 50 mins were measured. (**B**) In the first phase of the model, PSAM^4^-GlyR expressing mice exhibited significantly less nocifensive behaviours compared to mCherry expressing mice (mCherry: n = 6 mice, PSAM^4^-GlyR: n = 6 mice, unpaired t-test, * P < 0.05). Data expressed as mean ± S.E.M.

**Fig. S7.**
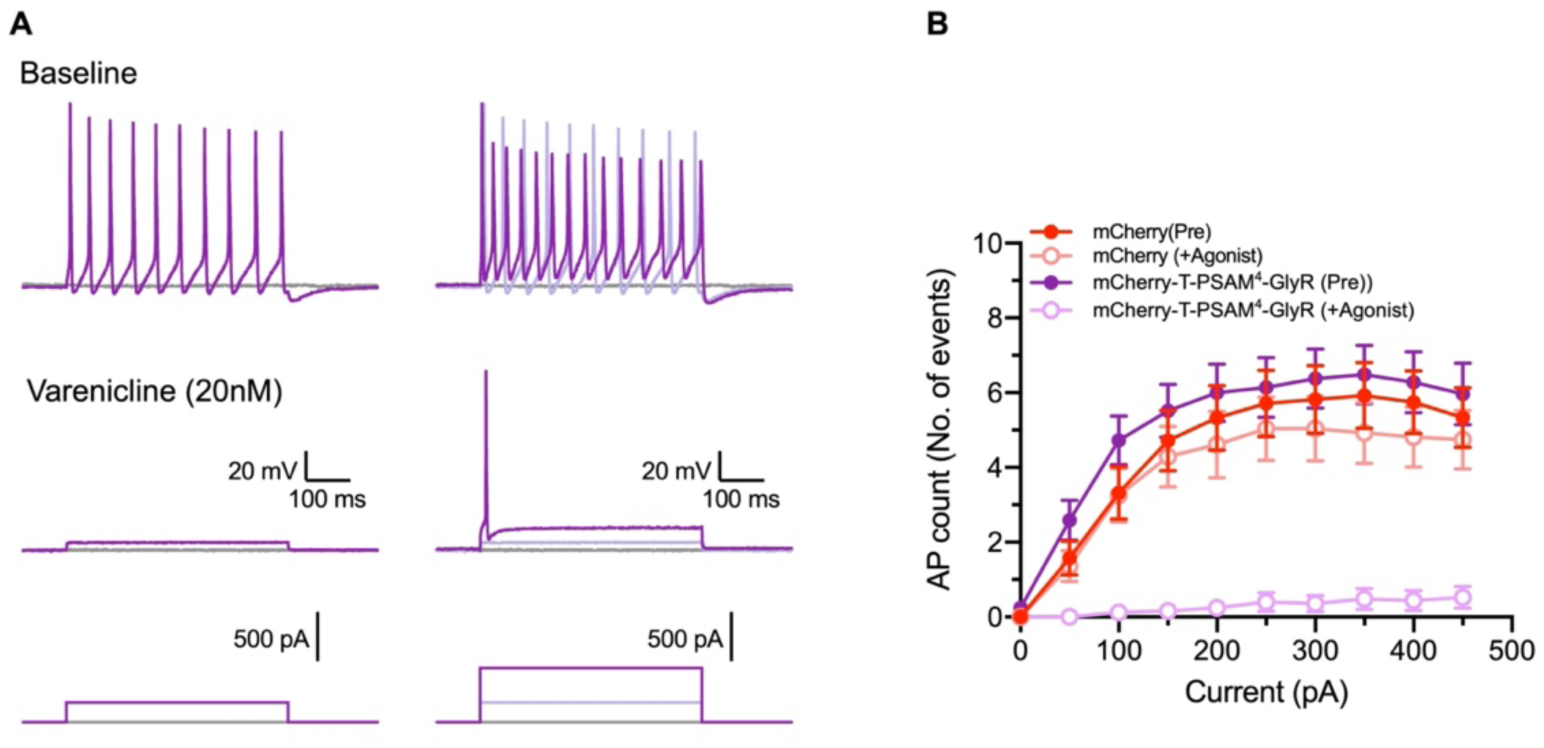
Repetitive firing of human-derived sensory neurons is abolished by PSAM^4^-GlyR activation. **(A)** Example traces showing repetitive firing with increasing current application before activation of PSAM^4^-GlyR (Top trace). After varenicline application, repetitive firing in PSAM^4^-GlyR expressing neurons was abolished. Neurons fired a single AP upon application of a large current pulse (Bottom trace). **(B)** Summary of quantification of the number of AP with increasing current pulses. Recordings in the presence of uPSEM^792^ and varenicline were pooled and labeled as Agonists. Data expressed as mean ± S.E.M.

## References

1. H. Breivik, B. Collett, V. Ventafridda, R. Cohen, D. Gallacher, Survey of chronic pain in Europe: prevalence, impact on daily life, and treatment. Eur. J. Pain 10, 287–333 (2006).

2. L. Colloca, T. Ludman, D. Bouhassira, R. Baron, A. H. Dickenson, D. Yarnitsky, R. Freeman, A. Truini, N. Attal, N. B. Finnerup, C. Eccleston, E. Kalso, D. L. Bennett, R. H. Dworkin, S. N. Raja, Neuropathic pain. Nat Rev Dis Primers 3, 17002 (2017).

3. D. A. Walsh, D. F. McWilliams, Mechanisms, impact and management of pain in rheumatoid arthritis. Nature Reviews Rheumatology 10, 581–592 (2014).

4. N. B. Finnerup, N. Attal, S. Haroutounian, E. McNicol, R. Baron, R. H. Dworkin, I. Gilron, M. Haanpaa, P. Hansson, T. S. Jensen, P. R. Kamerman, K. Lund, A. Moore, S. N. Raja, A. S. Rice, M. Rowbotham, E. Sena, P. Siddall, B. H. Smith, M. Wallace, Pharmacotherapy for neuropathic pain in adults: a systematic review and meta-analysis. Lancet Neurol. 14, 162–173 (2015).

5. S. Chakrabarti, L. A. Pattison, K. Singhal, J. R. F. Hockley, G. Callejo, E. S. J. Smith, Acute inflammation sensitizes knee-innervating sensory neurons and decreases mouse digging behavior in a TRPV1-dependent manner. Neuropharmacology 143, 49–62 (2018).

6. T. J. Boucher, S. B. McMahon, Neurotrophic factors and neuropathic pain. Curr. Opin. Pharmacol. 1, 66–72 (2001).

7. M. S. Gold, G. F. Gebhart, Nociceptor sensitization in pain pathogenesis. Nat. Med. 16, 1248–1257 (2010).

8. S. Haroutounian, L. Nikolajsen, T. F. Bendtsen, N. B. Finnerup, A. D. Kristensen, J. B. Hasselstrom, T. S. Jensen, Primary afferent input critical for maintaining spontaneous pain in peripheral neuropathy. Pain 155, 1272–1279 (2014).

9. K. C. Kajander, S. Wakisaka, G. J. Bennett, Spontaneous discharge originates in the dorsal root ganglion at the onset of a painful peripheral neuropathy in the rat. Neurosci. Lett. 138, 225–228 (1992).

10. R. G. Xie, D. W. Zheng, J. L. Xing, X. J. Zhang, Y. Song, Y. B. Xie, F. Kuang, H. Dong, S. W. You, H. Xu, S. J. Hu, Blockade of persistent sodium currents contributes to the riluzole-induced inhibition of spontaneous activity and oscillations in injured DRG neurons. PLoS One 6, e18681 (2011).

11. Y. A. Rho, S. A. Prescott, Identification of molecular pathologies sufficient to cause neuropathic excitability in primary somatosensory afferents using dynamical systems theory. PLoS Comput. Biol. 8, e1002524 (2012).

12. R. H. Gracely, S. A. Lynch, G. J. Bennett, Painful neuropathy: altered central processing maintained dynamically by peripheral input. Pain 51, 175–194 (1992).

13. S. G. Waxman, The molecular pathophysiology of pain: abnormal expression of sodium channel genes and its contributions to hyperexcitability of primary sensory neurons. Pain **Suppl** 6, S133–S140 (1999).

14. Y. Song, H. M. Li, R. G. Xie, Z. F. Yue, X. J. Song, S. J. Hu, J. L. Xing, Evoked bursting in injured Abeta dorsal root ganglion neurons: a mechanism underlying tactile allodynia. Pain 153, 657–665 (2012).

15. J. L. Xing, S. J. Hu, K. P. Long, Subthreshold membrane potential oscillations of type A neurons in injured DRG. Brain Res. 901, 128–136 (2001).

16. S. G. Waxman, G. W. Zamponi, Regulating excitability of peripheral afferents: emerging ion channel targets. Nat. Neurosci. 17, 153–163 (2014).

17. H. Starobova, A. Alshammari, I. G. Winkler, I. Vetter, The role of the neuronal microenvironment in sensory function and pain pathophysiology. J. Neurochem. **n/a**, (2022).

18. M. P. Klinck, J. S. Mogil, M. Moreau, B. D. X. Lascelles, P. A. Flecknell, T. Poitte, E. Troncy, Translational pain assessment: could natural animal models be the missing link? Pain 158, 1633–1646 (2017).

19. M. Alsaloum, G. P. Higerd, P. R. Effraim, S. G. Waxman, Status of peripheral sodium channel blockers for non-addictive pain treatment. Nat. Rev. Neurol. 16, 689–705 (2020).

20. P. Creamer, M. Hunt, P. Dieppe, Pain mechanisms in osteoarthritis of the knee: effect of intraarticular anesthetic. J. Rheumatol. 23, 1031–1036 (1996).

21. D. T. Demant, K. Lund, N. B. Finnerup, J. Vollert, C. Maier, M. S. Segerdahl, T. S. Jensen, S. H. Sindrup, Pain relief with lidocaine 5% patch in localized peripheral neuropathic pain in relation to pain phenotype: a randomised, double-blind, and placebo-controlled, phenotype panel study. Pain 156, 2234–2244 (2015).

22. A. Vaso, H. M. Adahan, A. Gjika, S. Zahaj, T. Zhurda, G. Vyshka, M. Devor, Peripheral nervous system origin of phantom limb pain. Pain 155, 1384–1391 (2014).

23. D. Atasoy, S. M. Sternson, Chemogenetic Tools for Causal Cellular and Neuronal Biology. Physiol. Rev. 98, 391–418 (2018).

24. S. Chakrabarti, L. A. Pattison, B. Doleschall, R. H. Rickman, H. Blake, G. Callejo, P. A. Heppenstall, E. S. J. Smith, Intraarticular Adeno-Associated Virus Serotype AAV-PHP.S-Mediated Chemogenetic Targeting of Knee-Innervating Dorsal Root Ganglion Neurons Alleviates Inflammatory Pain in Mice. Arthritis Rheumatol 72, 1749–1758 (2020).

25. R. E. Miller, S. Ishihara, B. Bhattacharyya, A. Delaney, D. M. Menichella, R. J. Miller, A. M. Malfait, Chemogenetic Inhibition of Pain Neurons in a Mouse Model of Osteoarthritis. Arthritis Rheumatol 69, 1429–1439 (2017).

26. J. L. Saloman, N. N. Scheff, L. M. Snyder, S. E. Ross, B. M. Davis, M. S. Gold, Gi-DREADD Expression in Peripheral Nerves Produces Ligand-Dependent Analgesia, as well as Ligand-Independent Functional Changes in Sensory Neurons. J Neurosci 36, 10769–10781 (2016).

27. S. M. Iyer, S. Vesuna, C. Ramakrishnan, K. Huynh, S. Young, A. Berndt, S. Y. Lee, C. J. Gorini, K. Deisseroth, S. L. Delp, Optogenetic and chemogenetic strategies for sustained inhibition of pain. Sci. Rep. 6, 30570 (2016).

28. J. L. Gomez, J. Bonaventura, W. Lesniak, W. B. Mathews, P. Sysa-Shah, L. A. Rodriguez, R. J. Ellis, C. T. Richie, B. K. Harvey, R. F. Dannals, M. G. Pomper, A. Bonci, M. Michaelides, Chemogenetics revealed: DREADD occupancy and activation via converted clozapine. Science 357, 503–507 (2017).

29. D. F. Manvich, K. A. Webster, S. L. Foster, M. S. Farrell, J. C. Ritchie, J. H. Porter, D. Weinshenker, The DREADD agonist clozapine N-oxide (CNO) is reverse-metabolized to clozapine and produces clozapine-like interoceptive stimulus effects in rats and mice. Sci. Rep. 8, 3840 (2018).

30. C. J. Magnus, P. H. Lee, D. Atasoy, H. H. Su, L. L. Looger, S. M. Sternson, Chemical and genetic engineering of selective ion channel-ligand interactions. Science 333, 1292–1296 (2011).

31. C. J. Magnus, P. H. Lee, J. Bonaventura, R. Zemla, J. L. Gomez, M. H. Ramirez, X. Hu, A. Galvan, J. Basu, M. Michaelides, S. M. Sternson, Ultrapotent chemogenetics for research and potential clinical applications. Science 364, (2019).

32. S. C. Gantz, M. M. Ortiz, A. J. Belilos, K. Moussawi, Excitation of medium spiny neurons by ’inhibitory’ ultrapotent chemogenetics via shifts in chloride reversal potential. Elife 10, (2021).

33. Z. Wu, H. Yang, P. Colosi, Effect of genome size on AAV vector packaging. Mol. Ther. 18, 80–86 (2010).

34. J. Y. Dong, P. D. Fan, R. A. Frizzell, Quantitative analysis of the packaging capacity of recombinant adeno-associated virus. Hum. Gene Ther. 7, 2101–2112 (1996).

35. K. W. Sung, M. Kirby, M. P. McDonald, D. M. Lovinger, E. Delpire, Abnormal GABA(A) receptor-mediated currents in dorsal root ganglion neurons isolated from Na-K-2Cl cotransporter null mice. J. Neurosci. 20, 7531–7538 (2000).

36. L. Batti, M. Mukhtarov, E. Audero, A. Ivanov, R. C. Paolicelli, S. Zurborg, C. Gross, P. Bregestovski, P. A. Heppenstall, Transgenic mouse lines for non-invasive ratiometric monitoring of intracellular chloride. Front. Mol. Neurosci. 6, 11 (2013).

37. K. I. Chisholm, N. Khovanov, D. M. Lopes, F. La Russa, S. B. McMahon, Large Scale In Vivo Recording of Sensory Neuron Activity with GCaMP6. eNeuro 5, (2018).

38. G. A. Weir, S. J. Middleton, A. J. Clark, T. Daniel, N. Khovanov, S. B. McMahon, D. L. Bennett, Using an engineered glutamate-gated chloride channel to silence sensory neurons and treat neuropathic pain at the source. Brain 140, 2570–2585 (2017).

39. L. He, W. Zhao, L. Zhang, M. Ilango, N. Zhao, L. Yang, Z. Guan, Modified Spared Nerve Injury Surgery Model of Neuropathic Pain in Mice. J Vis Exp, (2022).

40. R. Y. North, Y. Li, P. Ray, L. D. Rhines, C. E. Tatsui, G. Rao, C. A. Johansson, H. Zhang, Y. H. Kim, B. Zhang, G. Dussor, T. H. Kim, T. J. Price, P. M. Dougherty, Electrophysiological and transcriptomic correlates of neuropathic pain in human dorsal root ganglion neurons. Brain 142, 1215–1226 (2019).

41. J. Serra, H. Bostock, R. Sola, J. Aleu, E. Garcia, B. Cokic, X. Navarro, C. Quiles, Microneurographic identification of spontaneous activity in C-nociceptors in neuropathic pain states in humans and rats. Pain 153, 42–55 (2012).

42. D. L. Bennett, A. J. Clark, J. Huang, S. G. Waxman, S. D. Dib-Hajj, The Role of Voltage-Gated Sodium Channels in Pain Signaling. Physiol. Rev. 99, 1079–1151 (2019).

43. L. Cao, A. McDonnell, A. Nitzsche, A. Alexandrou, P. P. Saintot, A. J. Loucif, A. R. Brown, G. Young, M. Mis, A. Randall, S. G. Waxman, P. Stanley, S. Kirby, S. Tarabar, A. Gutteridge, R. Butt, R. M. McKernan, P. Whiting, Z. Ali, J. Bilsland, E. B. Stevens, Pharmacological reversal of a pain phenotype in iPSC-derived sensory neurons and patients with inherited erythromelalgia. Sci. Transl. Med. 8, 335ra356 (2016).

44. T. C. Alich, P. Roderer, B. Szalontai, K. Golcuk, S. Tariq, M. Peitz, O. Brustle, I. Mody, Bringing to light the physiological and pathological firing patterns of human induced pluripotent stem cell-derived neurons using optical recordings. Front. Cell. Neurosci. 16, 1039957 (2022).

45. J. Perez-Sanchez, Y. De Koninck, in *Oxford Research Encyclopedia of Neuroscience*. (2019).

46. B. Graham, S. Redman, A simulation of action potentials in synaptic boutons during presynaptic inhibition. J. Neurophysiol. 71, 538–549 (1994).

47. W. D. Willis, John Eccles’ studies of spinal cord presynaptic inhibition. Prog. Neurobiol. 78, 189–214 (2006).

48. R. Bardoni, T. Takazawa, C. K. Tong, P. Choudhury, G. Scherrer, A. B. Macdermott, Pre- and postsynaptic inhibitory control in the spinal cord dorsal horn. Ann. N. Y. Acad. Sci. 1279, 90–96 (2013).

49. W. D. Willis, Jr., Dorsal root potentials and dorsal root reflexes: a double-edged sword. Exp. Brain Res. 124, 395–421 (1999).

50. N. Doyon, L. Vinay, S. A. Prescott, Y. De Koninck, Chloride Regulation: A Dynamic Equilibrium Crucial for Synaptic Inhibition. Neuron 89, 1157–1172 (2016).

51. S. A. Hewitt, J. I. Wamsteeker, E. U. Kurz, J. S. Bains, Altered chloride homeostasis removes synaptic inhibitory constraint of the stress axis. Nat. Neurosci. 12, 438–443 (2009).

52. F. Ferrini, J. Perez-Sanchez, S. Ferland, L. E. Lorenzo, A. G. Godin, I. Plasencia-Fernandez, M. Cottet, A. Castonguay, F. Wang, C. Salio, N. Doyon, A. Merighi, Y. De Koninck, Differential chloride homeostasis in the spinal dorsal horn locally shapes synaptic metaplasticity and modality-specific sensitization. Nat Commun 11, 3935 (2020).

53. P. Takkala, Y. Zhu, S. A. Prescott, Combined Changes in Chloride Regulation and Neuronal Excitability Enable Primary Afferent Depolarization to Elicit Spiking without Compromising its Inhibitory Effects. PLoS Comput. Biol. 12, e1005215 (2016).

54. S. J. Middleton, A. M. Barry, M. Comini, Y. Li, P. R. Ray, S. Shiers, A. C. Themistocleous, M. L. Uhelski, X. Yang, P. M. Dougherty, T. J. Price, D. L. Bennett, Studying human nociceptors: from fundamentals to clinic. Brain 144, 1312–1335 (2021).

55. S. D. Dib-Hajj, A. M. Rush, T. R. Cummins, F. M. Hisama, S. Novella, L. Tyrrell, L. Marshall, S. G. Waxman, Gain-of-function mutation in Nav1.7 in familial erythromelalgia induces bursting of sensory neurons. Brain 128, 1847–1854 (2005).

56. M. Alsaloum, J. I. R. Labau, S. Liu, P. R. Effraim, S. G. Waxman, Stem cell-derived sensory neurons modelling inherited erythromelalgia: normalization of excitability. Brain 146, 359–371 (2023).

57. S. Ratté, S. A. Prescott, Afferent hyperexcitability in neuropathic pain and the inconvenient truth about its degeneracy. Curr. Opin. Neurobiol. 36, 31–37 (2016).

58. J. Raper, R. D. Morrison, J. S. Daniels, L. Howell, J. Bachevalier, T. Wichmann, A. Galvan, Metabolism and Distribution of Clozapine-N-oxide: Implications for Nonhuman Primate Chemogenetics. ACS Chem. Neurosci. 8, 1570–1576 (2017).

59. M. C. Walker, D. M. Kullmann, Optogenetic and chemogenetic therapies for epilepsy. Neuropharmacology 168, 107751 (2020).

60. C. Jimenez-Ruiz, I. Berlin, T. Hering, Varenicline: a novel pharmacotherapy for smoking cessation. Drugs 69, 1319–1338 (2009).

61. H. M. Faessel, R. S. Obach, H. Rollema, P. Ravva, K. E. Williams, A. H. Burstein, A review of the clinical pharmacokinetics and pharmacodynamics of varenicline for smoking cessation. Clin. Pharmacokinet. 49, 799–816 (2010).

62. D. Sharon, A. Kamen, Advancements in the design and scalable production of viral gene transfer vectors. Biotechnol Bioeng 115, 25–40 (2018).

63. N. Bessis, F. J. GarciaCozar, M. C. Boissier, Immune responses to gene therapy vectors: influence on vector function and effector mechanisms. Gene Ther. 11 **Suppl 1**, S10–17 (2004).

64. M. A. Kotterman, T. W. Chalberg, D. V. Schaffer, Viral Vectors for Gene Therapy: Translational and Clinical Outlook. Annu. Rev. Biomed. Eng. 17, 63–89 (2015).

65. F. Wang, Z. Qin, H. Lu, S. He, J. Luo, C. Jin, X. Song, Clinical translation of gene medicine. J. Gene Med. 21, e3108 (2019).

66. S. L. Ginn, A. K. Amaya, I. E. Alexander, M. Edelstein, M. R. Abedi, Gene therapy clinical trials worldwide to 2017: An update. J. Gene Med. 20, e3015 (2018).

67. R. S. Finkel, E. Mercuri, B. T. Darras, A. M. Connolly, N. L. Kuntz, J. Kirschner, C. A. Chiriboga, K. Saito, L. Servais, E. Tizzano, H. Topaloglu, M. Tulinius, J. Montes, A. M. Glanzman, K. Bishop, Z. J. Zhong, S. Gheuens, C. F. Bennett, E. Schneider, W. Farwell, D. C. De Vivo, Nusinersen versus Sham Control in Infantile-Onset Spinal Muscular Atrophy. New England Journal of Medicine 377, 1723–1732 (2017).

68. K. Y. Chan, M. J. Jang, B. B. Yoo, A. Greenbaum, N. Ravi, W. L. Wu, L. Sanchez-Guardado, C. Lois, S. K. Mazmanian, B. E. Deverman, V. Gradinaru, Engineered AAVs for efficient noninvasive gene delivery to the central and peripheral nervous systems. Nat. Neurosci. 20, 1172–1179 (2017).

69. B. E. Deverman, P. L. Pravdo, B. P. Simpson, S. R. Kumar, K. Y. Chan, A. Banerjee, W. L. Wu, B. Yang, N. Huber, S. P. Pasca, V. Gradinaru, Cre-dependent selection yields AAV variants for widespread gene transfer to the adult brain. Nat. Biotechnol. 34, 204–209 (2016).

70. X. Chen, S. Ravindra Kumar, C. D. Adams, D. Yang, T. Wang, D. A. Wolfe, C. M. Arokiaraj, V. Ngo, L. J. Campos, J. A. Griffiths, T. Ichiki, S. K. Mazmanian, P. B. Osborne, J. R. Keast, C. T. Miller, A. S. Fox, I. M. Chiu, V. Gradinaru, Engineered AAVs for non-invasive gene delivery to rodent and non-human primate nervous systems. Neuron 110, 2242–2257 e2246 (2022).

71. R. Dhandapani, C. M. Arokiaraj, F. J. Taberner, P. Pacifico, S. Raja, L. Nocchi, C. Portulano, F. Franciosa, M. Maffei, A. F. Hussain, F. de Castro Reis, L. Reymond, E. Perlas, S. Garcovich, S. Barth, K. Johnsson, S. G. Lechner, P. A. Heppenstall, Control of mechanical pain hypersensitivity in mice through ligand-targeted photoablation of TrkB-positive sensory neurons. Nat. Commun. 9, 1640 (2018).

72. Z.-Z. Xu, Y. H. Kim, S. Bang, Y. Zhang, T. Berta, F. Wang, S. B. Oh, R.-R. Ji, Inhibition of mechanical allodynia in neuropathic pain by TLR5-mediated A-fiber blockade. Nat. Med. 21, 1326–1331 (2015).

73. L. E. Fenno, J. Mattis, C. Ramakrishnan, M. Hyun, S. Y. Lee, M. He, J. Tucciarone, A. Selimbeyoglu, A. Berndt, L. Grosenick, K. A. Zalocusky, H. Bernstein, H. Swanson, C. Perry, I. Diester, F. M. Boyce, C. E. Bass, R. Neve, Z. J. Huang, K. Deisseroth, Targeting cells with single vectors using multiple-feature Boolean logic. Nat. Methods 11, 763–772 (2014).

74. Y. Liu, S. Hegarty, C. Winter, F. Wang, Z. He, Viral vectors for neuronal cell type-specific visualization and manipulations. Curr. Opin. Neurobiol. 63, 67–76 (2020).

75. J. Juttner, A. Szabo, B. Gross-Scherf, R. K. Morikawa, S. B. Rompani, P. Hantz, T. Szikra, F. Esposti, C. S. Cowan, A. Bharioke, C. P. Patino-Alvarez, O. Keles, A. Kusnyerik, T. Azoulay, D. Hartl, A. R. Krebs, D. Schubeler, R. I. Hajdu, A. Lukats, J. Nemeth, Z. Z. Nagy, K. C. Wu, R. H. Wu, L. Xiang, X. L. Fang, Z. B. Jin, D. Goldblum, P. W. Hasler, H. P. N. Scholl, J. Krol, B. Roska, Targeting neuronal and glial cell types with synthetic promoter AAVs in mice, non-human primates and humans. Nat. Neurosci. 22, 1345–1356 (2019).

76. D. H. Bryant, A. Bashir, S. Sinai, N. K. Jain, P. J. Ogden, P. F. Riley, G. M. Church, L. J. Colwell, E. D. Kelsic, Deep diversification of an AAV capsid protein by machine learning. Nat. Biotechnol. 39, 691–696 (2021).

77. S. Mouchbahani-Constance, C. Lagard, J. Schweizer, I. Labonté, M. Georgiopoulos, C. Otis, M. St-Louis, E. Troncy, P. Sarret, A. Ribeiro-Da-Silva, J. A. Ouellet, P. Séguéla, M.-E. Paquet, R. Sharif-Naeini, Modulating the activity of human nociceptors with a SCN10A promoter-specific viral vector tool. Neurobiology of Pain 13, 100120 (2023).

78. N. Percie du Sert, V. Hurst, A. Ahluwalia, S. Alam, M. T. Avey, M. Baker, W. J. Browne, A. Clark, I. C. Cuthill, U. Dirnagl, M. Emerson, P. Garner, S. T. Holgate, D. W. Howells, N. A. Karp, S. E. Lazic, K. Lidster, C. J. MacCallum, M. Macleod, E. J. Pearl, O. H. Petersen, F. Rawle, P. Reynolds, K. Rooney, E. S. Sena, S. D. Silberberg, T. Steckler, H. Wurbel, The ARRIVE guidelines 2.0: Updated guidelines for reporting animal research. PLoS Biol. 18, e3000410 (2020).

79. Z. Liu, O. Chen, J. B. J. Wall, M. Zheng, Y. Zhou, L. Wang, H. R. Vaseghi, L. Qian, J. Liu, Systematic comparison of 2A peptides for cloning multi-genes in a polycistronic vector. Sci. Rep. 7, 2193 (2017).

80. A. Arcourt, L. Gorham, R. Dhandapani, V. Prato, F. J. Taberner, H. Wende, V. Gangadharan, C. Birchmeier, P. A. Heppenstall, S. G. Lechner, Touch Receptor-Derived Sensory Information Alleviates Acute Pain Signaling and Fine-Tunes Nociceptive Reflex Coordination. Neuron 93, 179–193 (2017).

81. R. J. Carter, J. Morton, S. B. Dunnett, Motor coordination and balance in rodents. Curr. Protoc. Neurosci. **Chapter** 8, Unit 8 12 (2001).

